# Urban sensory conditions alter rival interactions and mate choice in urban and forest túngara frogs

**DOI:** 10.1101/2024.01.13.575508

**Authors:** Judith A.H. Smit, Vera Thijssen, Andrew D. Cronin, Jacintha Ellers, Wouter Halfwerk

## Abstract

Sexual communication often takes place in networks with multiple competing signalers being simultaneously assessed by mate choosers. Altered sensory conditions, such as noise and light pollution, can affect communication by altering signal production and perception. While evidence of sensory pollution affecting sexual signaling is widespread, few studies assess impacts on sexual signaling during rival interactions as well as mate choice, let alone whether urban and forest populations have diverged in their response. Here, we investigate the effects of urban sensory conditions on sexual communication in urban and forest túngara frogs (*Engystomops pustulosus*). We recorded dyadic vocal rival interactions and assessed mate choice with and without noise and light pollution in the lab. We show that urban sensory conditions can directly impact the intensity of rival interactions, differences between rivals and mate choice, though changes were often in opposite directions for frogs of urban and forest origins. Moreover, we demonstrate that changes in rival interactions in response to light and noise pollution can also indirectly affect how females choose between potential mates. Our study reveals origin-dependent direct and indirect effects of noise and light pollution, which ultimately translate into fitness consequences for sexual communication in urban populations.

## 1. Background

Sexual communication encompasses the production of and responses to sexual signals, such as song, pheromones, or visual displays, which have evolved in response to rival competition and mate choice (Darwin 1871; Andersson 1994). Most sexual signaling occurs in networks with multiple rivals and mate choosers, such as bird leks or frog choruses, characterized by spatiotemporal overlap in sexual signals (McGregor and Peake 2000; Gerhardt and Huber 2002). Specifically in the acoustic domain, signalers can often flexibly alter the timing and characteristics of sexual signals depending on their perception of nearby rivals (Otter et al. 2002; Greenfield 2005; Neelon and Höbel 2019; Larter et al. 2022), shaping temporal patterns and differences between signaling rivals (Greenfield 2015). Mate choosers evaluate rival interactions, and exert mate choice based on comparison between sexual signals (Mennill et al. 2002; Bateson and Healy 2005; Callander et al. 2013), often in a non-linear and irrational manner (Akre and Johnsen 2014; Lea and Ryan 2015). Since both sexual signaling and mate choice are shaped by phenotypes of conspecifics, considering the social environment is crucial for understanding sexual communication.

How sexual signalers and intended receivers, such as rivals and mates, interact strongly depends on the sensory environment, since sensory conditions can affect the production, transmission and perception of sexual signals (Wiley and Richards 1978; Brumm and Slabbekoorn 2005). This is particularly apparent in urban environments, where anthropogenic noise and artificial light at night (ALAN) are widespread (Barber et al. 2010; Kyba et al. 2017), and have been shown to impact sensory systems of many organisms (Halfwerk and Slabbekoorn 2015; Dominoni et al. 2020). Although animals can often alter their behavior in response to urban sensory conditions, either via plastic or evolutionary responses (Lowry et al. 2013; Sol et al. 2013), it is poorly understood how urban-induced behavioral changes in signalers and receivers influence sexual communication and ultimately fitness.

In the case of signalers, direct changes in the production of sexual signals (see Cronin, Smit, Muñoz, et al. 2022 for an extensive overview) can occur, for example, in response to noisy conditions by increasing the pitch or amplitude of their vocalizations (Cynx et al. 1998; Slabbekoorn and Peet 2003; Halfwerk et al. 2011; Lampe et al. 2012; Kunc et al. 2022), presumably to counter masking effects of noise (Halfwerk and Slabbekoorn 2015; Dominoni et al. 2020). Similarly, in response to light pollution animals can adjust their signal composition or timing (Kempenaers et al. 2010; van Geffen, Groot, et al. 2015). Moreover, anthropogenic noise can affect how an individual perceives and responds to a rival (Luther and Magnotti 2014; LaZerte et al. 2017), which presumably translates into altered rival interactions (e.g. changed temporal patterns or differences between signaling rivals) with potential consequences for mate choice. Most studies on signal adjustments under urban sensory conditions, however, take communication networks out of the equation by, for example, testing individuals in isolation or by using static playback designs (e.g. Smit et al. 2022). While such designs provide useful insights in direct effects of sensory pollutants on signalers under standardized social environments, they fail to assess how interactions between rivals and, consequently, mate choice are affected.

Urban sensory conditions can also directly impact mate choice (Candolin and Wong 2019), though the effects of sensory pollutants on mate detectability, discrimination and preference are far from clear (Halfwerk and Slabbekoorn 2015; Dominoni et al. 2020). Noise and light pollution generally seem to decrease mate attraction towards a sexual signal (Bird and Parker 2014; Huet des Aunay et al. 2014; van Geffen, van Eck, et al. 2015), most likely as a result of masking or misleading mechanisms (Halfwerk and Slabbekoorn 2015; Dominoni et al. 2020). Importantly, mating decisions are generally based on comparisons between sexual signals (Mennill et al. 2002; Bateson and Healy 2005; Callander et al. 2013), and sensory conditions can also affect how mate choosers evaluate and choose from multiple signals (Rand et al. 1997; Coss et al. 2020; Taylor et al. 2021). It is still poorly understood, however, how urban sensory conditions directly, as well as indirectly via altered rival interactions, affect mate choice between multiple signalers. Moreover, it remains to be clarified whether rivals and mate choosers from urban populations have changed their behaviors in response to urban sensory conditions.

The túngara frog (*Engystomops pustulosus*), commonly found in urban and forest areas, is an ideal study species to investigate the effects of altered sensory conditions on sexual communication in interactive settings, since males aggregate in choruses and compete using acoustic sexual signals on which females base their mating decisions (Ryan 1980; Ryan 1985). Calling behavior is strongly shaped by the social environment, as males increase their call complexity (by adding ‘chucks’, short high amplitude elements, to their calls), amplitude and call rate when their social environment becomes more competitive (Rand and Ryan 1981; Green 1990; Bernal et al. 2007; Halfwerk et al. 2016). Females generally prefer more conspicuous sexual signals (Rand and Ryan 1981; Bernal et al. 2009), and their preference is further shaped by rival differences and temporal dynamics (Akre et al. 2011; Tárano 2015; Larter and Ryan 2024). Frogs in urban areas produce more conspicuous calls than frogs in forest areas (Halfwerk et al. 2019), but not in the lab when using conspecific playbacks (Smit et al. 2022). Differences in the field seem partially related to urban sensory conditions (Cronin, Smit, and Halfwerk 2022; Smit et al. 2022). It is currently unclear, however, what the effects of sensory pollution are on how rivals interact and how females choose among them.

Our aim was to investigate how urban sensory conditions (noise and light pollution) impact rival interactions and mate choice, and whether the effects differ between urban and forest populations. We collected frogs from urban and forest field sites and recorded vocal interactions of rival pairs under urban and forest sensory conditions in our lab set-up. Subsequently, we played the recorded dyadic vocal interactions to females and quantified mate choice in phonotaxis experiments under urban or forest sensory conditions. For males, we predicted changes in rival interactions under noise and light pollution, since we know that sexual signaling in túngara frogs is affected by sensory conditions (Cronin, Smit, and Halfwerk 2022; Smit et al. 2022). Moreover, we predicted urban and forest males to be affected differently by urban sensory conditions, since it has been shown that behavioral as well as physiological responses to sensory pollutants depend on frog origin in túngara frog males (Halfwerk et al. 2019; Smit et al. 2024). Regarding mate choice, we predicted that urban sensory conditions would decrease female preference, because of the potentially masking, distracting, and misleading effects of noise and light pollution (Halfwerk and Slabbekoorn 2015; Dominoni et al. 2020), but we did not have expectations on whether urban and forest females would differ. Last, we predicted that altered rival differences in response to urban sensory pollution could have indirect consequences for mate choice.

## 2. Methods

### a) Study species and collection sites

We studied túngara frogs (*Engystomopos pustulosus*) from collection sites in the town of Gamboa and in Soberanía National Park, Republic of Panamá. We collected calling male frogs and amplectant pairs respectively 0-1 and 1-3 h after sunset from six urban and four forest sites that were minimally 190 m apart (Fig. S1, Tbl. S1). Urban sites were characterized by the occasional presence of humans, human-built structures, and light pollution (mean±sd, 1.28±1.10 lx, lux meter, HT Instruments HT309, Tbl. S1), whereas forest sites were located in forested areas free from artificial light at night (< 0.01 lx). We transported frogs in small plastic containers in a plastic cooler to facilities of the Smithsonian Tropical Research Institute (STRI) in Gamboa. We recorded male-male interactions between 19:00 and 2:30 in September 2022 and conducted phonotaxis experiments assessing female preference between 20:30 and 4:00 in November 2022. In total, we tested 120 males (forest: 48, urban: 72) of which 31 rival pairs (forest: 14, urban: 17) consisting of 41 males (forest: 17, urban: 24) interacted in both urban and forest sensory conditions and passed our criteria (see below). In the phonotaxis experiment, we tested 87 females (forest: 34, urban: 53, three were recaptured and retested), of which 80 females (forest: 31, urban: 49) made a total of 462 choices (forest: 170, urban: 292) out of 649 trials (see Tbl. S1 for details). After testing, we took mass (g), body size (SVL, mm) and ventral and dorsal photographs of all frogs to be able to identify recaptures. Pairs were released at the capture site on the same night, males were released three days after capture.

### b) Sensory treatments

We exposed urban and forest males and females to urban (urban noise and light) or forest (forest noise and light) sensory conditions using a full factorial design (Fig. 1). We played synthesized forest (∼50 dBA) or urban (∼70 dBA) noise (peak amplitude at frog, Voltcraft SL-100, fast, low, max; see 33,52 for details) using a speaker (Visaton FR8WP, frequency response 100 Hz to 20 kHz (−10 dB), connected to Renkforce T21 amplifiers) at ∼0.1 m from the male or two speakers at ∼1.7 m above the surface where females could move (Fig. S2). To ensure males and females would be exposed to similar noise levels, we calibrated the speakers daily using an artificial chorus playback (males: set to 67±2 dBA at 20 cm, females: set to 73±2 dBA at ∼1.7m; Voltcraft SL-100, fast, low, and max, Tbl. S2). Urban noise playbacks were mostly lower frequencies (<2 kHz), whereas the “whine” part of the túngara frog call range from ∼ 0.4 to 1 kHz and the energy in the “chuck” is concentrated >2 kHz, indicating some spectral overlap between the urban noise and the lowest frequency part of the call (for power spectra see Smit et al. 2022). To manipulate light conditions, we used white broad-spectrum LEDs (Nichia NSPW500DS, peak ∼460 nm) mimicking forest (<0.01 lx) or urban light levels (∼1.5 lx position of the male, ∼1.95 lx starting position of the female, HT instruments HT309, Fig. S2, Tbl. S2).

**Figure 1.**
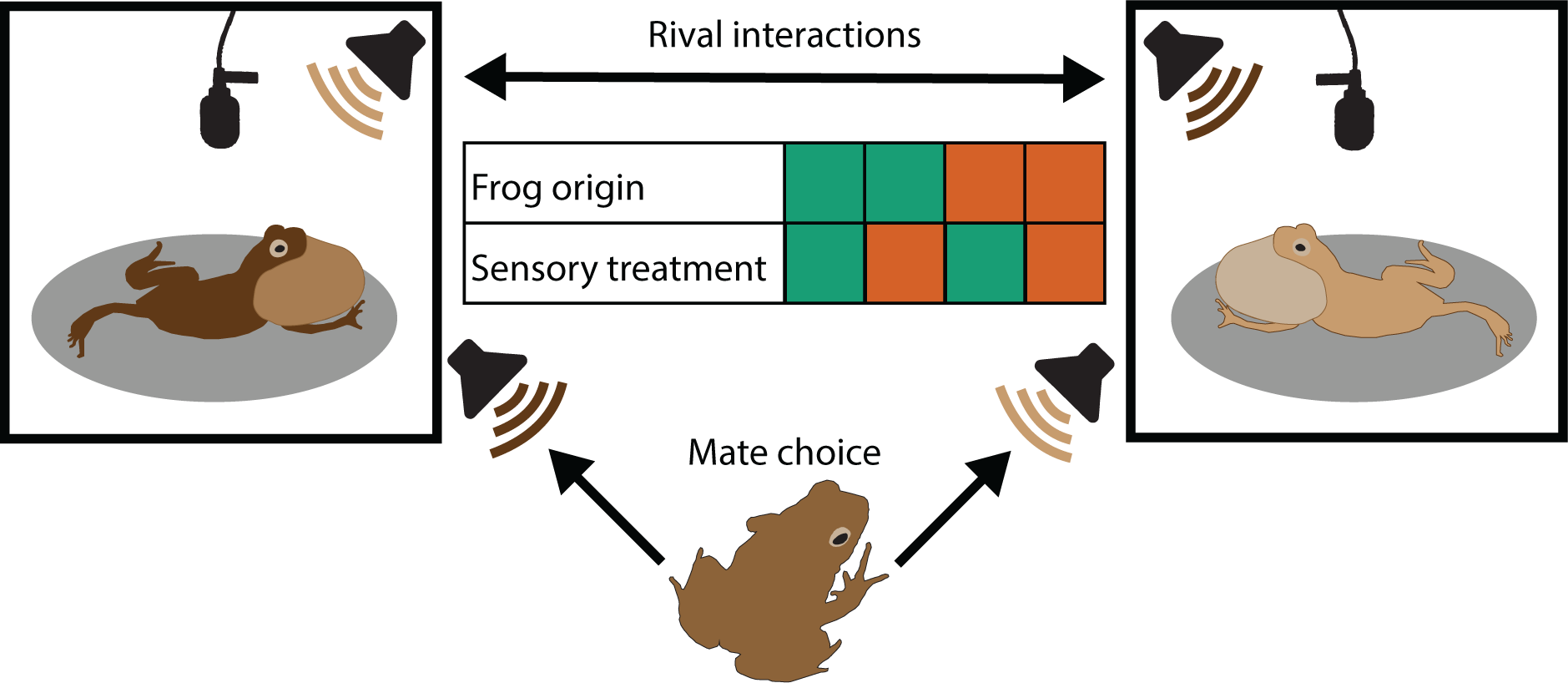
Schematic of experimental design. We recorded vocal interactions of rival pairs that could hear and respond to each other in acoustically connected recording boxes. The audio recordings of the rival interactions were later played to females in phonotaxis experiments to record mate choice. We conducted the experiments with rival pairs and females from urban and forest origins. During the experiments we exposed the frogs to urban or forest sensory treatments by manipulating light and noise levels in the recording boxes and phonotaxis chamber, using a full factorial design. Green and orange indicate, respectively, forest and urban origins and treatments.

### c) Recording male-male interactions

We recorded vocal interactions of males in two sets of four sound-attenuating recording boxes (L x W x H: 36 x 25 x 58 cm) under both forest and urban sensory conditions (Fig. 1). Our set-up allowed males to hear and respond to rivals by recording calls using omnidirectional lavalier microphones (AKG C417, 40 cm above the male) connected to two four-channel audio interfaces (Tascam US 4×4 HR and US 4×4) and live broadcasting the calls using the speakers that also played noise. Prior to testing each day, we played a 1.5 kHz tone (UE roll 2 Bluetooth speaker) in each recording box at the position of the frog and adjusted the microphone gain in this box so the tone would be equally broadcast (61+/− 1 dBA at 20 cm, Voltcraft SL-100, fast, low) in all connected boxes.

Upon arrival in the lab, males were placed in tanks (AltDesign, L x W x H: 18.4 x 29.2 x 12.7 cm) with damp paper towels and wild termites *ad libitum*. The next day each male was transferred into a small plastic container (9 cm ø, 5.5 cm height) containing mesh on the sides for the male to hold onto while calling, a pebble as shelter and saran wrap as an acoustically transparent lid. We filled the containers with ∼2.5 cm of dechlorinated water at least 0.5 h before the start of an experimental round using low level red light to reduce disturbance. Each round, eight males from the same collection site were randomly placed in the recording boxes (temperature 24 - 26 °C) without using light, where they were stimulated with a ∼67 dB artificial chorus for 0.5 h (6 x 3 min block) while acclimating to the first sensory treatment. Next, we turned on all microphones and waited per group of four acoustically connected boxes for two frogs to start calling before disconnecting microphones of the other two males. The two non-interacting males were not recorded and broadcast, but they were able to hear the calls of the two interacting males. After at least 10 min in which the rival pair could interact, all four microphones were turned on again until another pair of frogs started calling. To maximize the number of different rival pairs, we switched containers with frogs between the two groups of connected recording boxes without using light. This procedure was repeated for 1-2 h until all calling frogs had interacted with each other, and we started with the next group of males. We recorded three groups of eight males per night, which we repeated in the same order the following day in the second sensory treatment. In total, we tested five cohorts of frogs, with the order of the sensory treatments randomly assigned and observers blind to the identities of the frogs.

For the rival pairs that interacted with each other under both forest and urban conditions, we selected the longest interactions, excluding the first 2 min in which the rival pair could interact to minimize carry-over effects from previous interactions. An interaction was defined as a period of at least 20 sec that both males called with pauses lasting a maximum of 10 sec, including the call of the male that started and ended the interaction. We filtered (>300 and <4500 Hz, 24 dB roll-off) the resulting 35 rival pairs using Audacity (version 3.1.3; Audacity Team, 2021).

Next, we characterized the selected longest interactions in terms of interaction length (sec) and temporal overlap of rival calls (% of calls), and per frog call rate (calls / min), call complexity (chucks / call) and whine amplitude measured as both root mean square (RMS) and peak-to-peak (P2P) amplitude. To calculate amplitude, we selected up to three non-overlapping whines with at least 0.1 sec silence before and after the call. The analyzed calls were selected from the loudest calls (based on visual inspection of oscillograms) and were never chosen from the first or last three calls of the interaction, because those calls typically are less loud. We calculated amplitude using *seewave* v. 2.2.0 (Sueur et al. 2008) in R v. 4.2.2 (Team R Development Core 2022), while correcting for background noise by subtracting the average RMS or P2P of the background noise 0.1 to 0.02 sec before and 0.02 to 0.1 sec after the call. To correct for microphone differences, we used the RMS or P2P of a nightly recorded 1 kHz calibration tone (G.R.A.S. 42AB, 114 dB). We used linear amplitude values for statistical analyses, and dB scale for reporting estimates and visualizations. Last, we excluded four rival pairs that had very few (< 8) or low amplitude (< 60 dB whine RMS) calls in one of their interactions from further analyses, resulting in 31 rival pairs that interacted in both forest and urban sensory conditions.

### d) Phonotaxis experiments

For the phonotaxis experiments, we selected 15 out of 31 rival pairs by applying additional selection criteria on both their longest forest and urban interaction. These criteria included a minimum interaction length of 30 sec and no simultaneous silence of the frogs for more than 5 sec in the first or last 30 sec of the interaction or twice within 30 sec, to increase the likelihood that females would show phonotactic responses. Moreover, we excluded two rival pairs that had an interaction for more than 150 sec, to minimize the number of females making a mating decision before hearing a complete interaction. Using Audacity, we removed background noise between calls, added 10 sec of silence after the interactions and corrected amplitudes for microphone differences based on the RMS of the calibration tone. The final 15 rival pairs (forest: 6, urban: 9) consisted of 27 different males and resulted in 30 stimuli (recorded in forest or urban sensory conditions).

Phonotactic responses of female frogs to the rival interactions were assessed in a hemi-anechoic chamber (Acoustic Systems, Austin, TX, L x W x H: 2.7 × 1.8 × 1.78 m, Fig. S2 for set-up) under both urban and forest sensory conditions (Fig. 1). Upon arrival in the lab, pairs were placed in small plastic containers (9 cm ø, 5.5 cm height) with a damp paper towel on the bottom and saran wrap as a lid using low level red light. After at least 0.5 h in the lab, we transferred females to one of two sound-attenuating recording boxes in which females acclimatized for 0.5-1 h to the first sensory treatment. Next, without using additional light, we separated individual females from their mates and placed them in the phonotaxis chamber (temperature 24 - 27 °C) beneath an acoustically permeable funnel at 90 cm distance from two speakers (Sherman Oaks Orb Audio Mod1, connected to Audioengine N22 amplifiers) separated by a 70° angle. Each speaker played calls from one of the interacting males, with rival pair, male treatment and speaker (left or right) randomly assigned. Speaker volumes were calibrated daily using an artificial whine set to the P2P amplitude of the loudest whine (82±0.5 dBC) at 50 cm (Voltcraft SL-100, fast, low, and max).

We exposed each female to a rival interaction for 2 min before raising the funnel and scored her choice (defined as a speaker approach to within 10 cm for at least 3 sec) and decision latency (sec) using an infrared camera. We ceased the trial if a female did not move within the first 5 min after raising the funnel or any following 2 min, or if she did not exhibit phonotaxis within 10 min. Wall climbing or crossing the chamber back line opposite to the speakers was interpreted as a lack of motivation and scored as no choice. After four trials or after not making a choice twice, we returned the females to the containers with their mates using low level red light. The same acclimatization and testing procedures were repeated for the second sensory treatment. We alternated the order of the two sensory treatments between nights to prevent biases. In the forest and urban sensory treatments, we obtained, respectively, 152 and 140 choices made by urban females, and 90 and 80 choices made by forest females. In terms of choices for the different rival pairs, we obtained 186 choices for forest rival pairs (94 and 92 recorded in forest and urban conditions, respectively), and 276 choices for urban rival pairs (140 and 136 recorded in forest and urban conditions, respectively).

### e) Statistical analyses

We analyzed calling behavior and female preference with (generalized) linear mixed models using *lme4* v. 1.1-32 (Bates et al. 2015) in R v. 4.2.2. We assessed statistical significance of interactions between sensory treatment and frog origin and, in case of a non-significant interaction, their main effects origin using likelihood ratio tests. Alternatively, if the interaction term was found to be statistically significant, we tested treatment effects for urban and forest frogs using post-hoc tests to obtain estimates, standard errors, t ratios and p values using *emmeans* v. 1.8.5 (Lenth 2023). In case we found a trend in the interaction term (0.05 < p < 0.1), we report both main effects and post-hoc test results. All numerical covariates were standardized. We verified normality and absence of heteroskedasticity for all initial models and checked for overdispersion and zero-inflation for initial models with binomial distributions using *DHARMa* v. 0.4.6 (Hartig 2022). Graphs were created using raw data with *ggplot2* v. 3.4.1 (Wickham 2016). Complete model outputs can be found in the supplementary materials (Table S3-6).

To assess impacts of sensory treatment on the longest rival interactions, we ran models on maximum interaction length (sec) and overlap rate, and we calculated the average of two rivals and their absolute differences for each call behavior trait (call rate, complexity, whine amplitude as RMS and P2P). In addition, we calculated proportional rival differences by dividing absolute rival differences by the highest value of the two rivals according to Weber’s law (Weber 1948). In all calling behavior models, we included male treatment, male origin, and their interaction as fixed effects and rival pair and male collection site as random intercept effects. All models used Gaussian distributions (identity link function), except for models on call overlap rate, that had binomial distributions (logit link function) with the number of overlapping vs. non-overlapping calls as a response variable. To meet model assumptions, we log-transformed interaction length and sqrt-transformed absolute differences in RMS whine amplitude, call rate and complexity and proportional differences in call rate and complexity.

To examine sensory treatment effects on mate choice, we ran models on preference strength and latency to choose (sec). Preference strength (0-1) was defined as the deviation from equal choices for both rivals, with 0 indicating equal choices for each rival (e.g. 5 out of 10 choices for both rivals) and 1 indicating all choices for one rival (e.g. 10 out of 10 choices for one rival). In case we recorded an uneven number of choices for a stimulus or rival pair, we set equal choices between both rivals to the first possible number of choices above 50% (e.g. 2 out of 3 choices was set as a preference strength of 0 in case we recorded 3 choices). Stimuli or rival pairs for which we only obtained one choice in an origin-treatment combination were excluded from the analyses. Models on preference strength had a binomial distribution (logit link) and combined as a response variable the number of recorded choices deviating from equal choices for both rivals and the maximum possible deviation from equal choices for both rivals. To test direct sensory treatment effects on urban and forest females, we included the interaction between female sensory treatment and female origin as fixed effect, and stimulus as a random intercept effect. To test indirect sensory treatment effects on female preference via altered rival interactions, we included the interaction between male sensory treatment and male origin as a fixed effect, and rival pair and male collection site as random intercept effects. Models on latency to choose additionally contained female or male collection site, for, respectively, models on direct and indirect treatment effects, and female ID as random intercept effects. For latency models we used a Gaussian distribution (identity link) and log-transformed the response variable to meet model assumptions.

To assess whether preference strength was higher than expected by random choice, we compared whether our measured preference strength was outside of 95% confidence intervals based on 1000 simulation generating random choices (Tbl. S7). Using unweighted or weighted mean preference strength (based on the number of choices per rival pair or stimulus) did not impact the results. Finally, we tested associations between mate choice and interaction characteristics, by again running models (with binomial distributions and logit links) on preference strength per stimulus. We added as fixed factors interaction length (log-transformed), overlap rate, and the proportional difference in call rate, complexity and whine amplitude (RMS and P2P) and as random intercept effect the rival pair. We checked for multicollinearity, and all variance inflation factors were < 1.7 (Zuur et al. 2010). Next, we tested significance of associations via an information theoretic approach using *MuMIn* v. 1.47.5 (Bartoń 2023), and we report weighted averaged model estimates and associated p values for models < 4 AIC (small sample size corrected) from the top model (Tbl. S8).

## Results

### Urban sensory conditions decrease maximum interaction length

Sensory treatment impacted maximum interaction length in an origin-dependent way (treatment x origin: n = 62, χ2 = 4.17, P = 0.041, Fig. 2A). The longest interactions between forest rivals were 51.0% shorter under urban compared to forest conditions (n = 28, t = 4.56, P < 0.001) and urban rivals showed a similar, but smaller, trend of 24.2% decrease (n = 34, t = 1.98, P = 0.056). How often two rivals overlapped their calls in these interactions did not depend on sensory treatment across or within origin (all P > 0.27, Tbl. S3, Fig. S3A).

**Figure 2.**
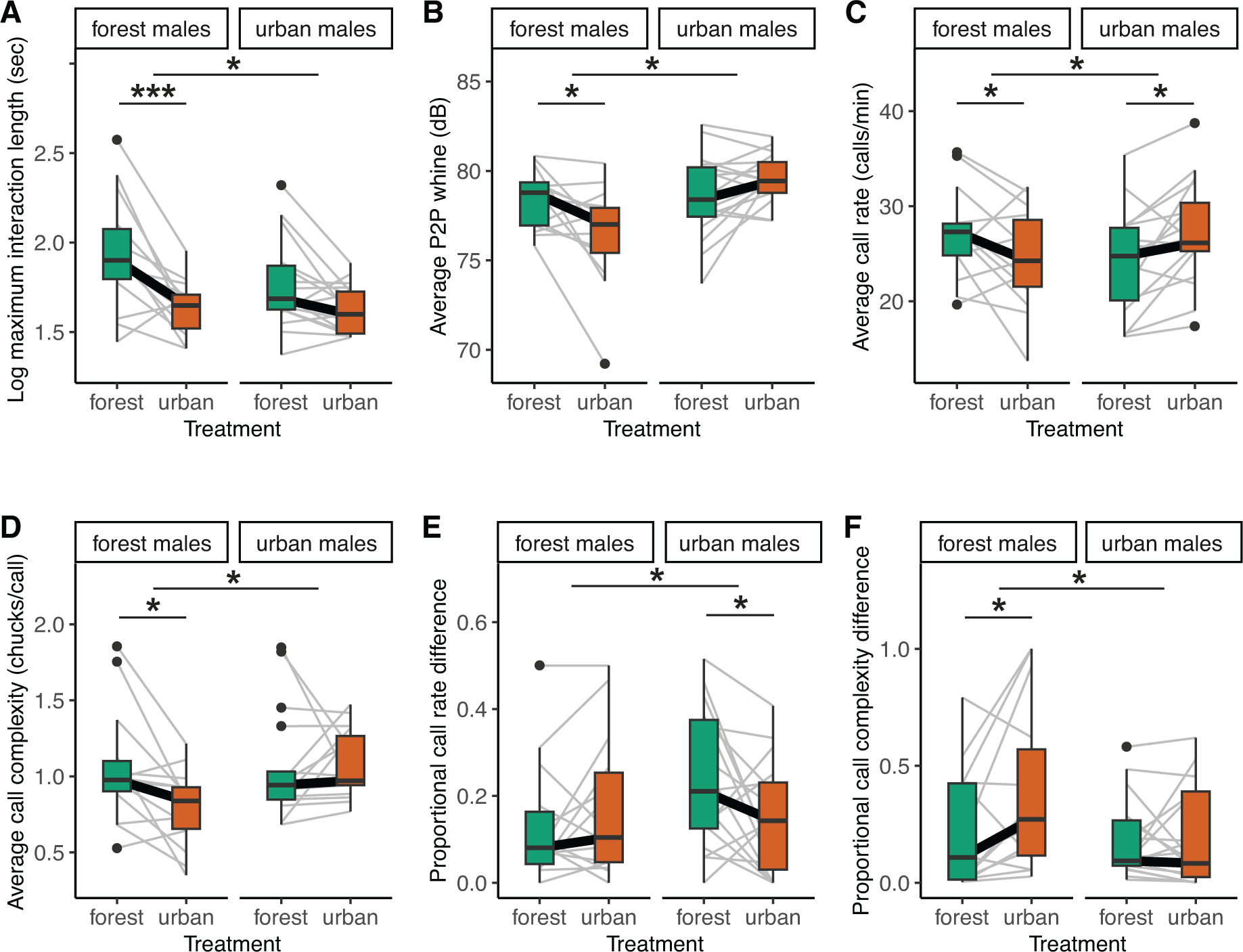
Effects of urban and forest sensory treatment on vocal interactions in urban and forest rival pairs. A) Maximum interaction length (log sec). Averages of two rivals in a pair for B) peak-to-peak (P2P) whine amplitude (dB), C) call rate (calls/min) and D) call complexity (chucks/call). Proportional differences (absolute difference divided by highest value) between two rivals in E) call rate and F) call complexity. Graphs show raw data and grey lines indicate rival pairs. Asterisks indicate statistically significance (* P < 0.05, *** P < 0.001) of interaction effects between treatment and origin, and of treatment effects within origins, see main text and Tbl. S3-5 for statistics.

### Increased intensity in rival interactions in familiar sensory conditions

When examining the average calling behavior of rival pairs during their longest interaction, we found the effect of sensory treatment to depend on male origin for whine amplitude (treatment x origin: n = 62, P2P: χ2 = 8.60, P = 0.003, Fig. 2B; RMS: χ2 = 5.77, P = 0.016, Fig. S3B), call rate (χ2 = 10.49, P = 0.001, Fig. 2C) and complexity (χ2 = 5.34, P = 0.021, Fig. 2D). Forest males decreased their whine amplitude (n = 28, P2P: −1.6 dB, t = 2.68, P = 0.011; RMS: −1.5 dB, t = 2.60, P = 0.014), call rate (−9.6%, t = 2.09, P = 0.045) and complexity (−25.2%, t = 2.86, P = 0.008) in urban compared to forest conditions. Urban males, on the other hand, increased their call rate when exposed to urban sensory conditions (+13.2%, n = 34, t = −2.79, P = 0.009), but not their amplitude or complexity (all P > 0.11).

### Proportional rival differences change with sensory conditions

The absolute difference between two rivals in calling behavior during their longest interaction showed a trend in the interaction between sensory treatment and origin when looking at call rate (treatment x origin, n = 62, χ2 = 3.37, P = 0.07, Fig. S3E, Tbl. S4) but did not reveal an overall treatment effect (χ2 = 1.44, P = 0.23). For urban rivals, absolute differences in call rate decreased when exposed to urban sensory conditions (−48.5%, n = 34, t = 2.14, P = 0.04), but no pattern was found for forest rivals (n = 28, t = −0.52, P = 0.61). Absolute differences in other aspects of calling behavior between two rivals were not affected by sensory treatment (all P > 0.24, Fig. S3C-F, Tbl. S4). When scaling absolute differences to the highest value of the rivals (proportional differences), we report interaction effects between sensory treatment and origin for call rate (χ2 = 4.57, P = 0.033, Fig. 1E, Tbl. S5) and complexity (χ2 = 6.16, P = 0.013, Fig. 1F). Urban males decreased their proportional differences in call rate in urban conditions (−0.11, t = 2.45, P = 0.020), while forest males increased their proportional differences in call complexity in urban conditions (+0.17, t = −2.86, P = 0.007). We found no effects of sensory treatment on proportional rival differences in call amplitude (all P > 0.43; Fig. S3G-H).

### Urban conditions directly affect female preference

The strength of female preference (i.e. how much preference deviated from equal choices for both rivals), was affected by sensory treatment in an origin-dependent way (treatment x origin, n = 462, χ2 = 6.66, P = 0.010, Fig. 3A, Tbl. S6). The urban sensory treatment increased female preference in urban females (n = 292, z ratio = −2.75, P = 0.006), but did not significantly alter preference in forest females (n = 170, z ratio = 1.19, P = 0.236). When testing whether preference strength deviated from random choices (Fig. S5, Tbl. S7), urban females showed a preference strength that was significantly higher than random ([0.55, 95% CI random choices [0.10; 0.42]) in urban conditions, but not in forest conditions (0.34, [0.11; 0.40]). Forest females showed the opposite pattern, with a significantly higher preference compared to random choices under forest conditions (0.61, [0.08; 0.48]), but not under urban conditions (0.42, [0.10; 0.50]). Last, latency to choose did not differ depending on sensory treatment (all P > 0.63, Fig. S4, Tbl. S6).

**Figure 3.**
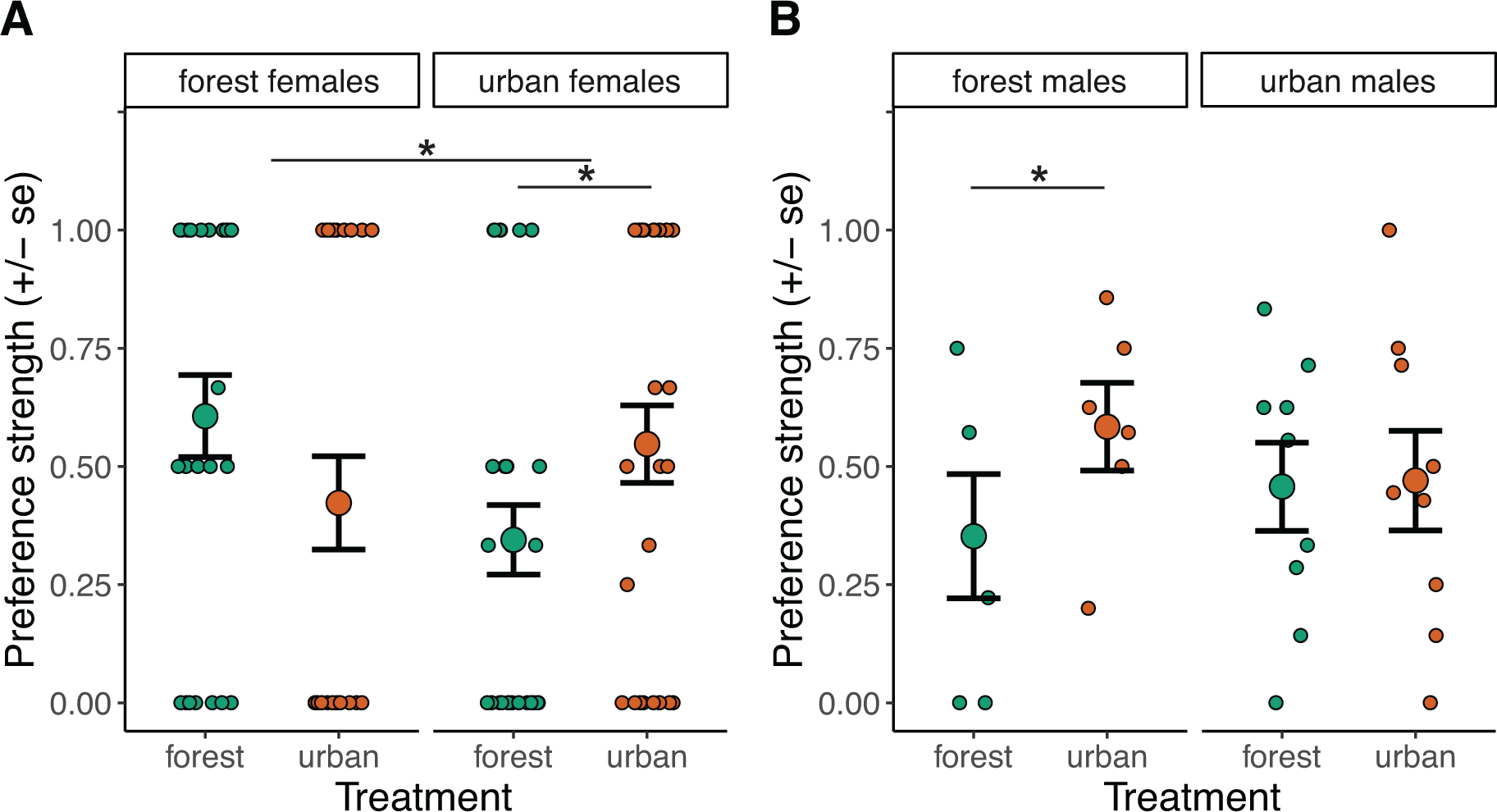
Mean (± standard error) preference strength (0 = no preference between two males, 1 = all choices for one male). Data is split in A) preference of urban and forest females choosing under urban or forest sensory conditions and in B) preference for urban and forest rival pairs interacting under urban or forest sensory conditions. Data points depict preference strength per stimulus (panel A) or per rival pair (panel B). Graphs show raw data. Asterisks indicate statistically significance (* P < 0.05) of interaction effects between treatment and origin, and of treatment effects within origins, see main text and Tbl. S7 for statistics.

### Urban-induced changes in forest rival interactions impact mate choice

We tested whether alterations in rival interactions due to urban sensory conditions affect female preference strength (Fig. 3B, Tbl. S6) and detected a trend in an interaction effect between male origin and treatment (treatment x origin: n = 462, χ2 = 2.89, P = 0.089, Fig 3B). While we found no overall treatment effect (χ2 = 1.97, P = 0.16), the urban sensory treatment increased preference strength when females chose from forest rivals (n = 186, z ratio = −2.17, P = 0.030), but not when choosing from urban rivals (n = 276, z ratio = −0.02, P = 0.985). When choosing from forest rival pairs interacting under forest conditions, females did not show a statistically significant preference (0.35, random choices 95% CI: [0.00; 0.38], Fig. S5, Tbl. S7), but when forest males interacted under urban conditions female preference was significantly higher than expected from random choices (0.58, [0.00; 0.39]). For urban rival pairs, females showed a statistically significant preference when choosing from urban rival pairs interacting under both forest (0.46, [0.03; 0.32]) and urban conditions (0.46, [0.04, 0.33]). Latency to choose did not differ depending on the sensory treatment in which the rivals interacted (all P > 0.19, Fig. S4, Tbl. S6). Overall, higher preference strength was significantly associated with higher proportional call rate difference (z = 2.48, P = 0.01, Tbl. S8) and lower interaction length (z = 2.30, P = 0.02), but not any other interaction level trait (all P > 0.25).

## 3. Discussion

Research on sensory pollution has often focused on individuals or communities, but rarely on interactions between individuals (Kurvers and Hölker 2015), which is particularly important for sexual communication, as this often takes place in networks with multiple simultaneous signalers and mate choosers (McGregor and Peake 2000; Gerhardt and Huber 2002). Our aim was therefore to assess effects of urban sensory conditions on both rival interactions and mate choice in urban and forest túngara frogs. We repeatedly recorded dyadic vocal interactions of rival pairs, with and without noise and light pollution, allowing us to quantify emergent interaction level properties (maximum interaction length, interaction intensity, overlap rate and rival differences), and subsequently tested female preference for the calls of the interacting males under similar sensory treatments. We found that urban sensory conditions affected both rival interactions and mate choice, with the direction of effects often depending on whether frogs were collected in urban or forest areas.

### Urban conditions affect urban and forest rival interactions differently

Maximum interaction length was significantly shorter under urban sensory conditions in forest rival pairs and showed the same trend in urban rival pairs. Since frogs are known to cease calling sooner under higher perceived predation risk (Tuttle et al. 1982), this pattern could be explained by urban noise being more disturbing and urban light being perceived as more risky because of increased visual detectability of the frogs (Rand et al. 1997). The stronger decrease in interaction length in forest rival pairs under noise and light pollution is in line with forest individuals being more vigilant (Halfwerk et al. 2019), a pattern also often reported for other taxa (Møller 2010; Uchida et al. 2019). Rival interactions can also be shortened when rivals perceive a clearer difference in the quality of their signals and therefore can settle their competition faster (Arnott and Elwood 2008), thereby reducing energetic and predation costs. This alternative explanation would suggest that the shorter interactions under urban conditions result from larger differences, as we found in forest but not urban rival pairs, or better perception of acoustic phenotypes of rivals, although we do not know of a perceptual mechanism supporting this last explanation. Furthermore, how often two rivals would overlap their calls was not affected by sensory treatment, which is in line with an earlier study with túngara frogs using static playbacks (Smit et al. 2022).

The intensity of the longest interaction between two rivals was impacted by urban sensory conditions, and, in line with our predictions, the direction of the effect depended on frog origin. Under urban sensory conditions the forest rival pairs lowered their average call rate, amplitude and complexity, whereas the urban rival pairs increased their call rate when exposed to urban conditions. Rivalling frogs thus had more intense interactions in sensory conditions matching the conditions of their origin, indicating that exposure to unfamiliar sensory conditions could have distracted or stressed the frogs leading to more conservative or de-escalating calling strategies (Goutte et al. 2010). Urban noise and light pollution have been shown to increase conspicuousness of sexual signaling in túngara frogs when exposed to a static playback (Smit et al. 2022), which is in line with our results on urban but not on forest rival interactions. Moreover, a translocation experiment between urban and forest sites indicated that only urban frogs show behavioral flexibility (Halfwerk et al. 2019), whereas our findings show that forest frogs also respond flexibly to sensory conditions. In forest areas, producing attractive sexual signals is balanced by higher predation and parasitic pressures (Ryan et al. 1982; Bernal et al. 2006; Akre et al. 2011), while predator and parasite abundances are much lower in urban areas or under urban sensory pollutants reducing the costs of producing conspicuous sexual signals (McMahon et al. 2017; Halfwerk et al. 2019). In terms of predation and parasitic costs, urban but not forest rival pairs thus seem to adjust their interaction intensity in an adaptive way to urban sensory pollutants, implying that urban males have adapted their behavior to urban sensory conditions.

In addition to the average calling behavior of a rival pair, it is insightful to examine differences between rivals’ acoustic phenotypes because rival differences often determine relative attractiveness to mate choosers. Absolute differences generally remained under urban conditions, indicating that rivals matched their calls to the same degree (e.g. Gerhardt et al. 2000), except for a smaller difference in call rate between urban rivals when exposed to urban sensory conditions. However, proportional rival differences (absolute difference divided by the highest value) are thought to be more important than absolute differences in mate choice (Weber 1948; Akre and Johnsen 2014). Urban sensory conditions lead to lower proportional call rate differences between urban rivals, and to higher proportional call complexity differences between forest rivals, possibly because of the opposite directions of the changes in interaction intensity in urban and forest rival pairs. Changes in absolute or proportional differences can have consequences for discriminating between rivals and therefore can affect mate choice, especially when potential mates are evaluated simultaneously like in túngara frogs (Bateson and Healy 2005; Akre et al. 2011; Callander et al. 2013).

### Effects of urban sensory conditions on mate choice

When perceiving and responding to sexual signals of potential mates, mate choice can directly be influenced by human-induced changes to the environment (Candolin and Wong 2019), but the effects of urban sensory pollutants on mate choice are currently understudied (Cronin, Smit, Muñoz, et al. 2022). Our study sheds light on female choice between sexual signals of two interacting rivals, quantified as preference strength (deviation from equal choice between the two rivals). We show that preference strength was directly affected by urban sensory conditions, but in an opposite way for urban and forest females. In line with our predictions, forest females showed a weaker (but not significantly lower) preference under urban sensory conditions, whereas to our surprise urban females increased their preference when exposed to noise and light pollution. Our findings indicate that preference strength is strongest in conditions matching the females’ origin, suggesting decreased attention to the sexual signals when making a choice under unfamiliar, possibly perceived as distracting or more risky, conditions (Bonachea and Ryan 2011). While variation in latency to choose could have provided insights in potential motivational or discriminatory differences between urban and forest females under the different sensory treatments, we did not find any effects on latency. Future research should investigate mechanisms underlying differences between urban and forest female mate choice behavior.

Generally, drastic environmental changes seem to directly affect mate choice resulting in lower sexual selection on signals because of interference with signal perception. Exposure to invasive plant chemicals can, for example, decrease female preference for males with stronger immune responses in Palmate newts (*Lissotriton helveticus*) (Iglesias-Carrasco et al. 2017). In our study, urban sensory conditions led to lower preference strength only in forest females, whereas urban females showed a higher preference strength under these conditions, suggesting they might have adapted to urban sensory conditions (Sol et al. 2013; Harding et al. 2019). Similarly, a recent study in bend-legged ground crickets (*Dianemobius nigrofasciatus*) has demonstrated that urban females reared in a common garden were faster in localizing an acoustic sexual signal under noisy conditions compared to forest females (Kuriwada 2023 but see Costello and Symes 2014)

In our female choice experiment, we also examined the indirect effects of urban sensory conditions on mate choice via urban-induced changes in rival interactions. We found that the sensory conditions in which the rivals interacted affected mate choice, but only for rival pairs from forest areas. For forest rivals, females had a higher preference strength when they chose from interactions recorded under urban sensory conditions compared to under forest sensory conditions. The higher preference strength for forest rival interactions recorded under urban conditions is in line with our finding that higher female preference strength was associated with shorter interactions, since forest rivals significantly decreased their maximum interaction length when exposed to noise and light pollution. Rival competitions can be settled earlier when rival differences are larger, making it possibly also easier for females to evaluate sexual signals. Indeed, we found an increase in proportional call complexity differences in forest rival pairs interacting in urban conditions, although we did not find a significant association with preference strength (but see Akre et al. 2011). For urban rival pairs we reported lower absolute and proportional call rate differences in interactions under urban conditions and we found a positive association between proportional call rate difference and female preference, but, surprisingly, we did not detect changes in preference strength depending on the recording conditions of urban rival pairs. Overall, we demonstrate that changes in how rivals interact when exposed to urban sensory conditions can indirectly affect mate choice, and thus have fitness consequences. We speculate that urban-induced changes in mate choice could have led to higher sexual selection on signaling, potentially driving the emergence of more conspicuous urban sexual signaling.

The current study focused on dyadic rival interactions with one female frog choosing at the time, the simplest form of a communication network. Communication networks can, however, consist out of many more signalers and mate choosers, resulting in complex chorus dynamics impacting female preference (Greenfield and Rand 2000; Calsbeek et al. 2022; Larter and Ryan 2024). Moreover, sensory conditions can affect communication networks more indirectly through changes in, for example, community composition, spatial distribution and species interactions (Francis et al. 2009; Naguib 2013). Our study can form a basis for understanding the effects of urbanization on sexual communication in interactive settings, and further work could build upon this by examining more complex communication networks.

### Conclusion

By examining dyadic rival interactions and letting females choose from playbacks of these interactions, we studied the effects of urbanization on a communication network in which sexual signaling as well as mate choice could be shaped by the phenotypes of conspecifics. Our study demonstrated that noise and light pollution can impact both signalers and, directly as well as indirectly, mate choosers. Interestingly, the effects of urban sensory pollution were often origin-dependent and urban and forest frogs showed opposite responses. When tested in sensory conditions matching the frogs’ origin, we found that rival interactions were most intense and preference strength was highest, suggesting that urban frogs might have adapted their sexual communication behavior, via plasticity and/or genetic processes, to urbanization.

## Acknowledgements

We would like to thank Argelis Sanchez for her help and dedication during the phonotaxis experiments. We are grateful to Ryan Taylor and Kim Hunter for the use of their phonotaxis chamber, and STRI for providing logistical support. Peter Moran and Matías Muñoz contributed with helpful discussions related to the data and the experimental set-up.

## Data availability

Analyses reported in this article can be reproduced using the R scripts and data provided by Smit (2023) via

## Supplementary material

**Figure S1.**
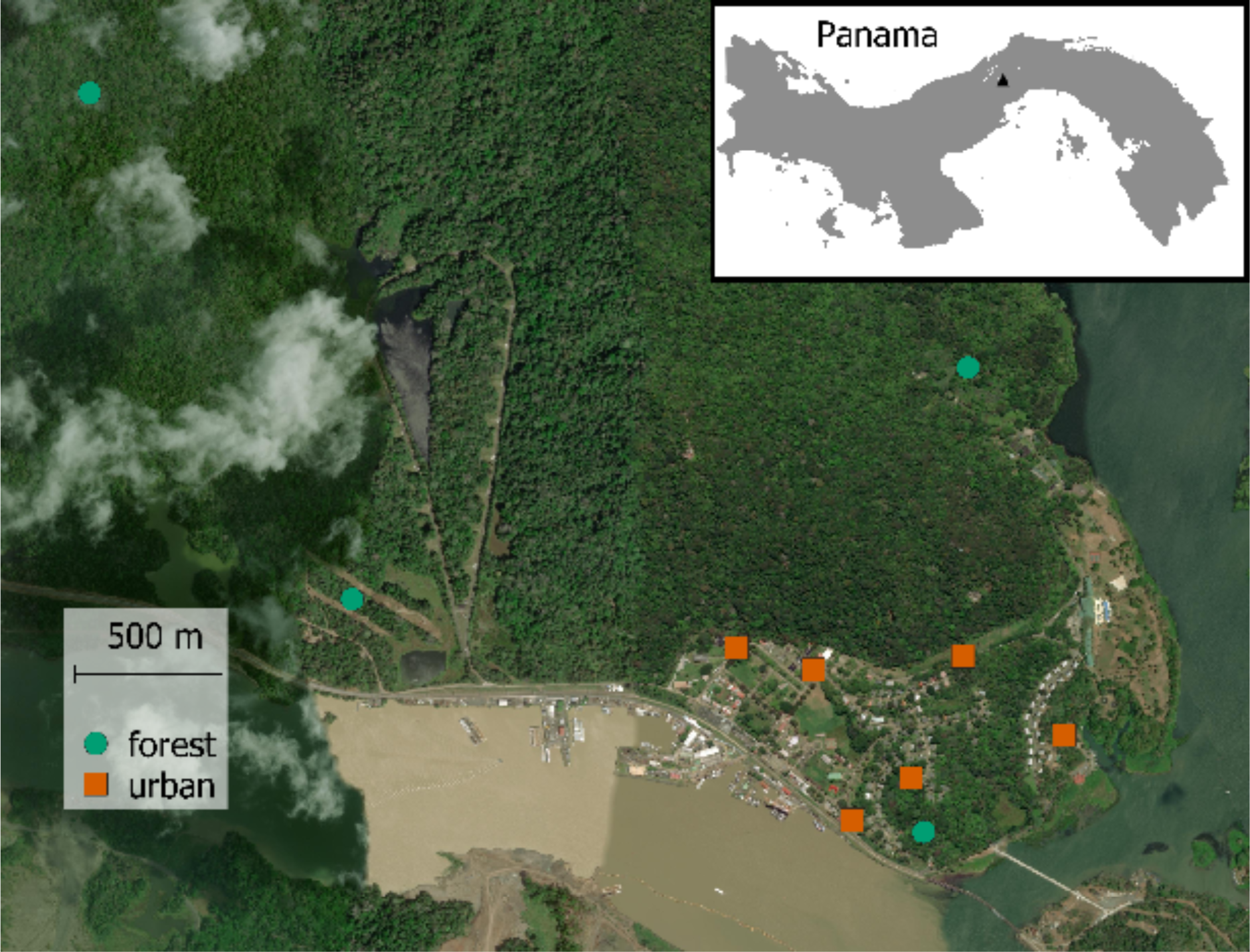
Map showing the locations of the 6 urban and 4 forest sites (see Tbl. S1 for coordinates and description)

**Table S1.** Overview of urban and forest frog collection sites. Mean and standard deviation of night-time light levels per site are based on five male calling locations. The number of collected males, the number of recorded interacting males and the subset used as phonotaxis stimuli are depicted per collection site. Similarly, the number of collected amplexed pairs, unique number of collected females and number of females that made one or more choices during the phonotaxis experiment are indicated per collection site.

| Site | Coordinates<br>(EPSG:4326 -<br>WGS84 CRS) | Mean±sd<br>light level<br>(lux) | Collected<br>males | Rival<br>interaction<br>males<br>(phonotaxis<br>stimuli) | Collected<br>pairs<br>(unique<br>females) | Choosing<br>females |
| --- | --- | --- | --- | --- | --- | --- |
| *Urban collection sites* |  |  |  |  |  |  |
| Stairs | 9.115101,<br>-79.699862 | 1.97±1.69 | 8 | 4 (3) | 12 (11) | 10 |
| Santa Cruz | 9.120339,<br>-79.703418 | 1.57±1.40 | 16 | 7 (3) | 11 (10) | 9 |
| Insectaries | 9.119689,<br>-79.701056 | 2.96±4.81 | 16 | 5 (5) | 19 (19) | 19 |
| 183 | 9.116392,<br>-79.698036 | 0.11±0.21 | 8 | 2 (0) | 3 (3) | 2 |
| Kent's<br>Marsh | 9.120090,<br>-79.696427 | 0.20±0.39 | 8 | 2 (0) | 1 (1) | 1 |
| Marina | 9.117693,<br>-79.693297 | 0.85±1.08 | 16 | 4 (4) | 9 (9) | 8 |
| *Forest collection sites* |  |  |  |  |  |  |
| Pipeline<br>Bridge | 9.137279,<br>-79.723399 | 0.00±0.00 | 16 | 6 (5) | 1 (1) | 0 |
| Pipeline<br>Start | 9.121832,<br>-79.715290 | 0.00±0.00 | 8 | 3 (2) | 7 (6) | 4 |
| Ditch | 9.114733,<br>-79.697640 | 0.00±0.00 | 8 | 3 (2) | 12 (12) | 12 |
| La Chunga | 9.128899,<br>-79.696274 | 0.00±0.00 | 16 | 5 (2) | 15 (15) | 15 |

**Figure S2.**
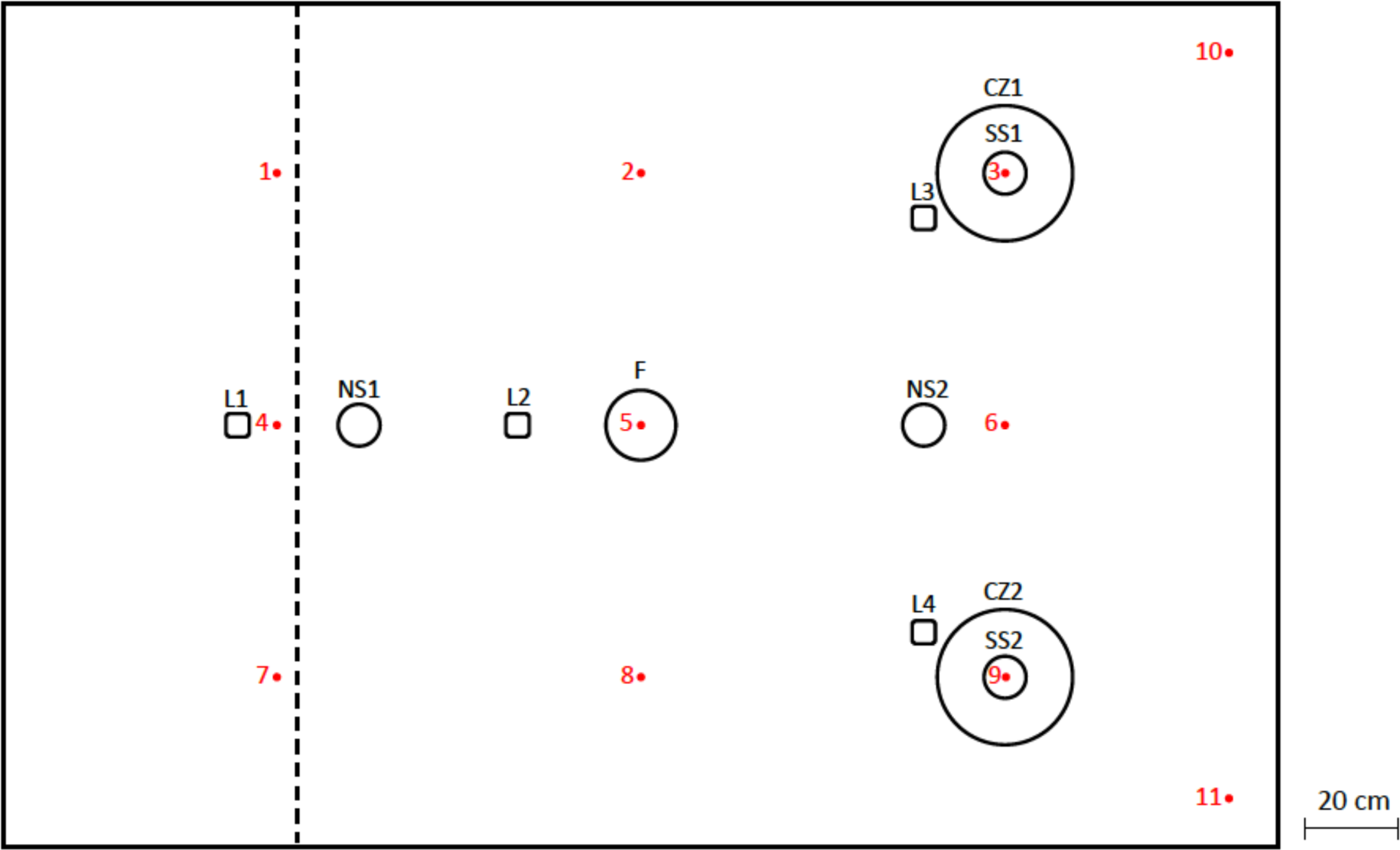
The set-up in the phonotaxis chamber on scale. L1-L4 = light 1-4; NS1-NS2 = noise speaker 1−2; F = funnel; SS1-SS2 = stimulus speaker 1−2; CZ1-CZ2 = choice zone of stimulus speaker 1−2; red dot 1-11 = calibration point 1-11. SS1 and SS2 are placed on the ground directed towards F, L1 and L2 are positioned on the ceiling in an angle so that their maximum light levels are right under NS1 and F and NS1, NS2, L3 and L4 are hanging on the ceiling facing downwards.

**Table S2.** Light and sound levels of the urban and forest light treatment, and of the chorus used to calibrate the noise levels at 11 calibration points in the phonotaxis chamber (see Fig. S2) measured on ground level with respectively HT instruments HT309 and Voltcraft SL-100 (A-weighted, fast, low, and max).

| Calibration Point | 1 | 2 | 3 | 4 | 5 | 6 | 7 | 8 | 9 | 10 | 11 |
| --- | --- | --- | --- | --- | --- | --- | --- | --- | --- | --- | --- |
| Light level urban (lx) | 0.07 | 0.15 | 1.08 | 0.98 | 1.95 | 0.27 | 0.07 | 0.11 | 1.16 | 0.00 | 0.00 |
| Light level forest (lx) | 0.00 | 0.00 | 0.00 | 0.00 | 0.00 | 0.00 | 0.00 | 0.00 | 0.00 | 0.00 | 0.00 |
| Chorus (dBA) | 69.0 | 71.7 | 68.6 | 71.1 | 73.7 | 71.1 | 68.2 | 71.9 | 69.1 | 65.9 | 65.7 |

**Table S3.**
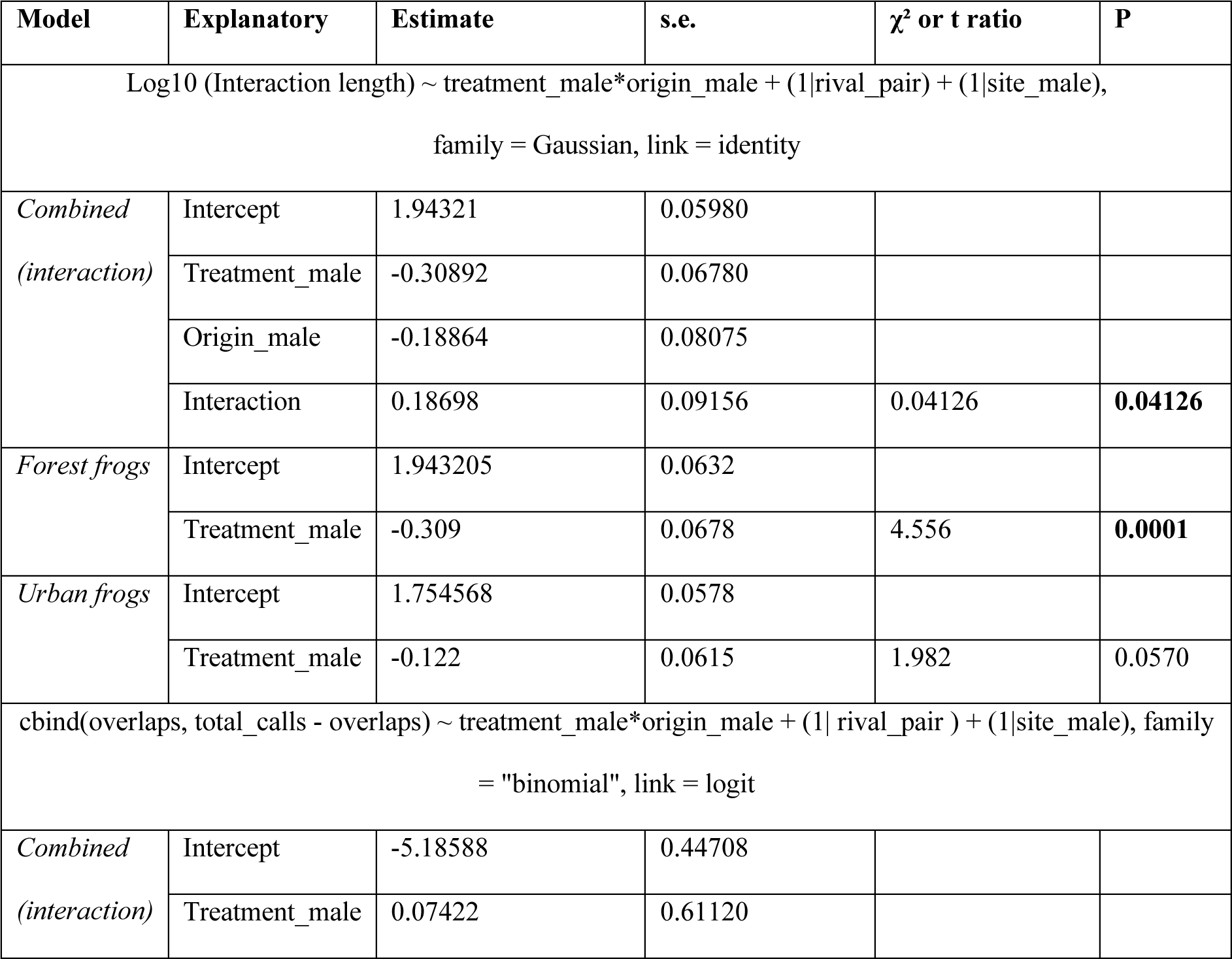
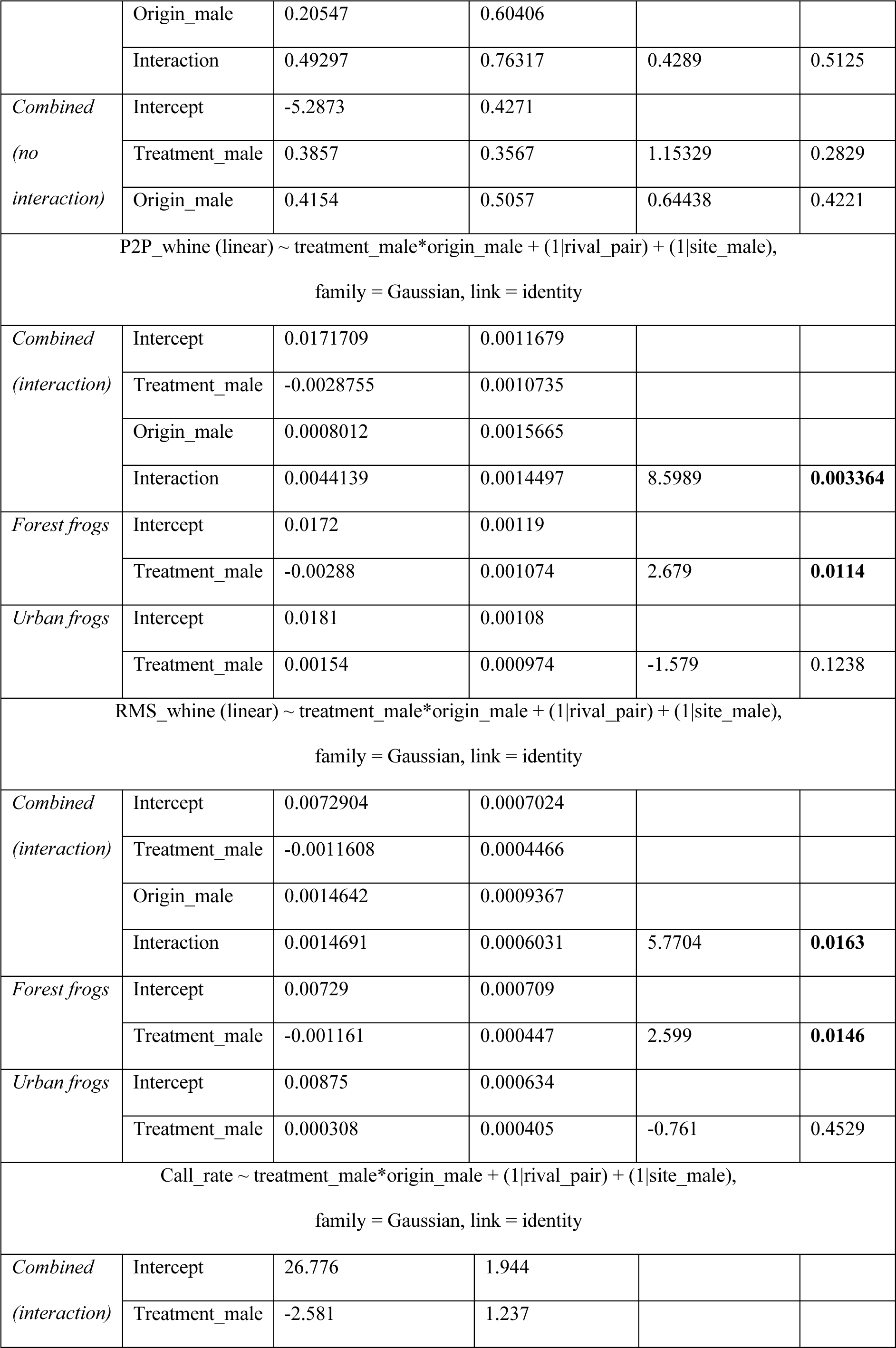
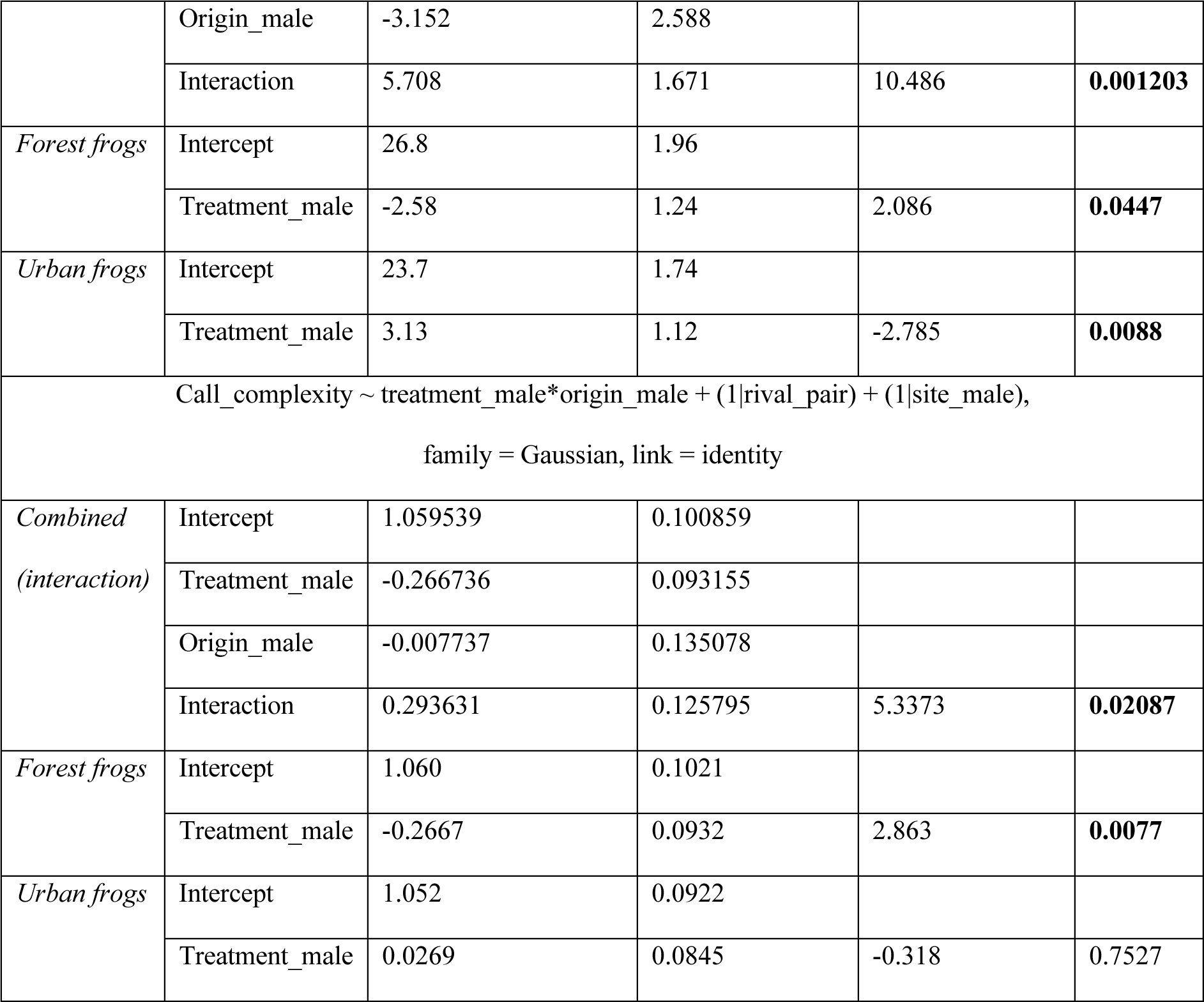
Results on the effects of male treatment and male origin on interaction level traits (maximum interaction length, overlap rate, whine amplitude (P2P and RMS), call rate and complexity). Initial models (GLMM) are specified below. χ² values and p values were obtained by likelihood ratio tests of models with and without the factor of interest, starting with the interaction term, and estimates and standard errors were obtained from the summary table of the model. In case of a non-significant interaction term, we subsequently removed the interaction term and report analyses testing for main effects of sensory treatment and frog origin. Alternatively, if the interaction team was found to be statistically significant, we tested treatment effects for urban and forest frogs using post-hoc tests to obtain estimates, standard errors, t ratios and p values. In case we found a trend in the interaction term (0.05 < p < 0.1), we report both main effects and post-hoc test results. Estimates for ‘treatment_male’ and ‘origin_male’ indicate the effects of urban compared to forest treatment or origin, estimates are not back transformed.

**Table S4.**
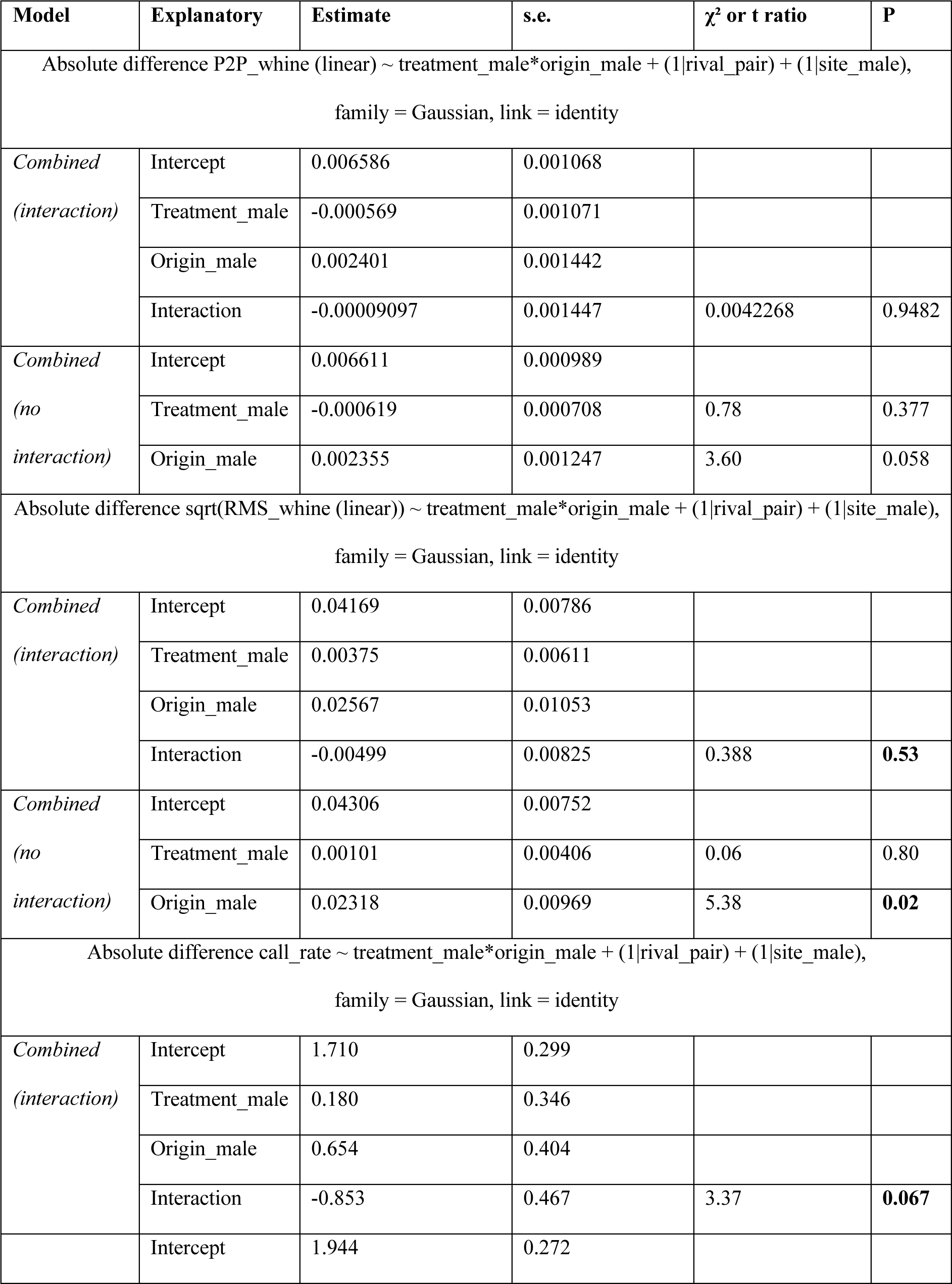
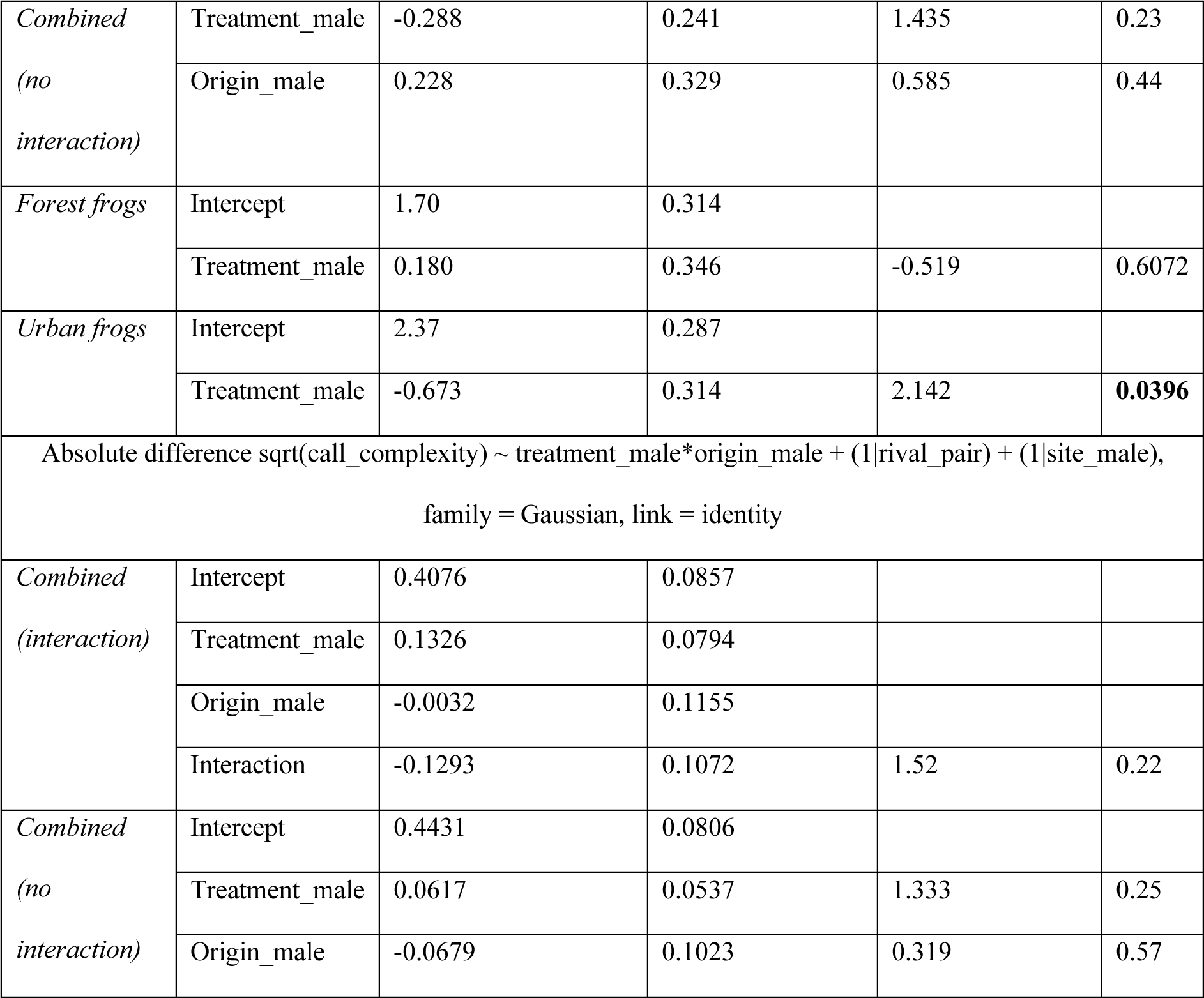
Results on the effects of male treatment and male origin on absolute rival differences (whine amplitude (P2P and RMS), call rate and complexity). See Table S3 for details.

| Model | Explanatory | Estimate | s.e. | $\chi^2$ or t ratio | P |
| --- | --- | --- | --- | --- | --- |
| Absolute difference P2P_whine (linear) ~ treatment_male*origin_male + (1\|rival_pair) + (1\|site_male),<br>family = Gaussian, link = identity |  |  |  |  |  |
| *Combined*<br>*(interaction)* | Intercept | 0.006586 | 0.001068 |  |  |
|  | Treatment_male | -0.000569 | 0.001071 |  |  |
|  | Origin_male | 0.002401 | 0.001442 |  |  |
|  | Interaction | -0.00009097 | 0.001447 | 0.0042268 | 0.9482 |
| *Combined*<br>*(no*<br>*interaction)* | Intercept | 0.006611 | 0.000989 |  |  |
|  | Treatment_male | -0.000619 | 0.000708 | 0.78 | 0.377 |
|  | Origin_male | 0.002355 | 0.001247 | 3.60 | 0.058 |
| Absolute difference sqrt(RMS_whine (linear)) ~ treatment_male*origin_male + (1\|rival_pair) + (1\|site_male),<br>family = Gaussian, link = identity |  |  |  |  |  |
| *Combined*<br>*(interaction)* | Intercept | 0.04169 | 0.00786 |  |  |
|  | Treatment_male | 0.00375 | 0.00611 |  |  |
|  | Origin_male | 0.02567 | 0.01053 |  |  |
|  | Interaction | -0.00499 | 0.00825 | 0.388 | **0.53** |
| *Combined*<br>*(no*<br>*interaction)* | Intercept | 0.04306 | 0.00752 |  |  |
|  | Treatment_male | 0.00101 | 0.00406 | 0.06 | 0.80 |
|  | Origin_male | 0.02318 | 0.00969 | 5.38 | **0.02** |
| Absolute difference call_rate ~ treatment_male*origin_male + (1\|rival_pair) + (1\|site_male),<br>family = Gaussian, link = identity |  |  |  |  |  |
| *Combined*<br>*(interaction)* | Intercept | 1.710 | 0.299 |  |  |
|  | Treatment_male | 0.180 | 0.346 |  |  |
|  | Origin_male | 0.654 | 0.404 |  |  |
|  | Interaction | -0.853 | 0.467 | 3.37 | **0.067** |
|  | Intercept | 1.944 | 0.272 |  |  |
| *Combined*<br>*(no interaction)* | Treatment_male | -0.288 | 0.241 | 1.435 | 0.23 |
|  | Origin_male | 0.228 | 0.329 | 0.585 | 0.44 |
| *Forest frogs* | Intercept | 1.70 | 0.314 |  |  |
|  | Treatment_male | 0.180 | 0.346 | -0.519 | 0.6072 |
| *Urban frogs* | Intercept | 2.37 | 0.287 |  |  |
|  | Treatment_male | -0.673 | 0.314 | 2.142 | **0.0396** |
| Absolute difference sqrt(call_complexity) ~ treatment_male*origin_male + (1\|rival_pair) + (1\|site_male),<br>family = Gaussian, link = identity |  |  |  |  |  |
| *Combined*<br>*(interaction)* | Intercept | 0.4076 | 0.0857 |  |  |
|  | Treatment_male | 0.1326 | 0.0794 |  |  |
|  | Origin_male | -0.0032 | 0.1155 |  |  |
|  | Interaction | -0.1293 | 0.1072 | 1.52 | 0.22 |
| *Combined*<br>*(no interaction)* | Intercept | 0.4431 | 0.0806 |  |  |
|  | Treatment_male | 0.0617 | 0.0537 | 1.333 | 0.25 |
|  | Origin_male | -0.0679 | 0.1023 | 0.319 | 0.57 |

**Table S5.** Results on the effects of male treatment and male origin on proportional rival differences (whine amplitude (P2P and RMS), call rate and complexity). See Table S3 for details.

| Model | Explanatory | Estimate | s.e. | $\chi^2$ or t ratio | P |
| --- | --- | --- | --- | --- | --- |
| Proportional difference P2P_whine (linear) ~ treatment_male*origin_male + (1\|rival_pair) + (1\|site_male),<br>family = Gaussian, link = identity |  |  |  |  |  |
| *Combined*<br>*(interaction)* | Intercept | 0.3136 | 0.0432 |  |  |
|  | Treatment_male | 0.0271 | 0.0400 |  |  |
|  | Origin_male | 0.0857 | 0.0583 |  |  |
|  | Interaction | -0.0878 | 0.0540 | 2.7 | 0.1 |
| *Combined*<br>*(no interaction)* | Intercept | 0.3376 | 0.0407 |  |  |
|  | Treatment_male | -0.0210 | 0.0276 | 0.592 | 0.44 |
|  | Origin_male | 0.0418 | 0.0517 | 0.692 | 0.41 |
| Proportional difference RMS_whine (linear) ~ treatment_male*origin_male + (1\|rival_pair) + (1\|site_male),<br>family = Gaussian, link = identity |  |  |  |  |  |
| *Combined*<br>*(interaction)* | Intercept | 0.2526 | 0.0594 |  |  |
|  | Treatment_male | 0.0609 | 0.0492 |  |  |
|  | Origin_male | 0.1795 | 0.0800 |  |  |
|  | Interaction | -0.0980 | 0.0664 | 2.24 | 0.13 |
| *Combined*<br>*(no interaction)* | Intercept | 0.27951 | 0.05658 |  |  |
|  | Treatment_male | 0.00719 | 0.03370 | 0.05 | 0.828 |
|  | Origin_male | 0.13053 | 0.07273 | 3.40 | 0.065 |
| Proportional difference call_rate ~ treatment_male*origin_male + (1\|rival_pair) + (1\|site_male),<br>family = Gaussian, link = identity |  |  |  |  |  |
| *Combined*<br>*(interaction)* | Intercept | 0.3303 | 0.0584 |  |  |
|  | Treatment_male | 0.0427 | 0.0636 |  |  |
|  | Origin_male | 0.1171 | 0.0786 |  |  |
|  | Interaction | -0.1842 | 0.0858 | 4.57 | **0.033** |
| *Forest frogs* | Intercept | 0.323 | 0.0602 |  |  |
|  | Treatment_male | 0.0427 | 0.0636 | -0.672 | 0.5064 |
| *Urban frogs* | Intercept | 0.452 | 0.0550 |  |  |
|  | Treatment_male | -0.1415 | 0.0577 | 2.454 | **0.0196** |
| Proportional difference call_complexity ~ treatment_male*origin_male + (1\|rival_pair) + (1\|site_male),<br>family = Gaussian, link = identity |  |  |  |  |  |
| *Combined*<br>*(interaction)* | Intercept | 0.38116 | 0.07406 |  |  |
|  | Treatment_male | 0.17807 | 0.06237 |  |  |
|  | Origin_male | -0.00579 | 0.09978 |  |  |
|  | Interaction | -0.21257 | 0.08422 | 6.16 | **0.013** |
| *Forest frogs* | Intercept | 0.381 | 0.0771 |  |  |
|  | Treatment_male | 0.1781 | 0.0624 | -2.855 | **0.0079** |
| *Urban frogs* | Intercept | 0.375 | 0.0706 |  |  |
|  | Treatment_male | -0.0345 | 0.0566 | 0.610 | 0.5469 |

**Figure S3.**
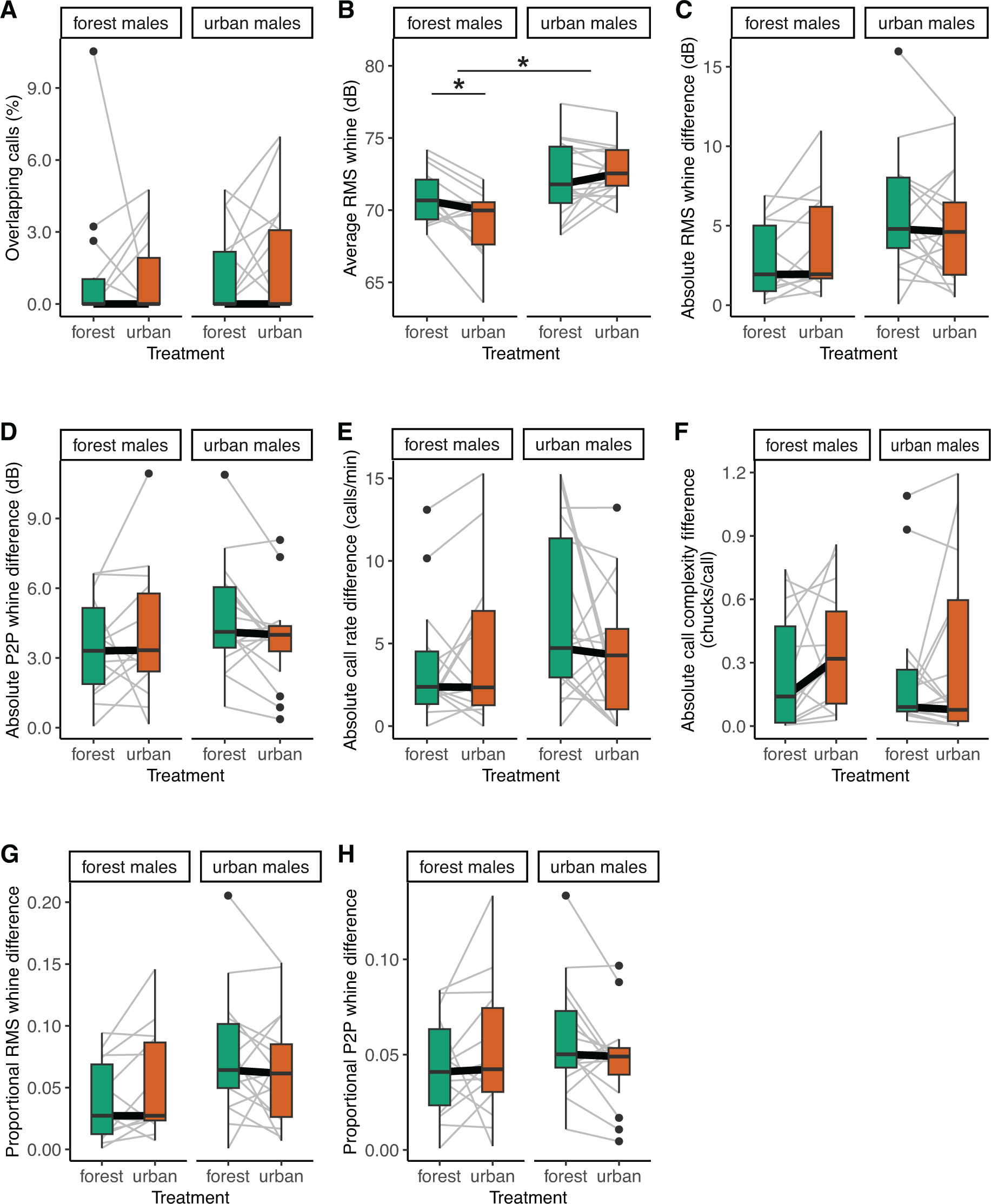
Effects of urban and forest sensory treatment on vocal interactions in urban and forest rival pairs. A) overlap rate (%), B) averages of rival pair in RMS whine amplitude (dB), absolute differences in C) RMS whine amplitude (dB), D) P2P whine amplitude, E) call rate and F) call complexity, proportional differences (absolute difference divided by highest value) between two rivals in G) RMS whine amplitude and H) P2P whine amplitude. Graphs show raw data, grey lines indicate rival pairs. Asterisks indicate statistically significance (* P < 0.05) of interaction effects between treatment and origin, and of treatment effects within origins, see main text and Tbl. S3-5 for statistics.

**Table S6.** Results on the effects of female/male treatment and female/male origin on latency to choose (seconds) and preference strength (0-1). See Table S3 for details on statistics.

| Model | Explanatory | Estimate | s.e. | $\chi^2$ or t ratio | P |
| --- | --- | --- | --- | --- | --- |
| Log10 (latency) ~ treatment_female*origin_female + (1\|female_ID) + (1\|stimulus) +<br>(1\|site_female), family = Gaussian, link = identity |  |  |  |  |  |
| *Combined*<br>*(interaction)* | Intercept | 1.8103 | 0.0465 |  |  |
|  | Treatment_female | -0.0234 | 0.0497 |  |  |
|  | Origin_female | -0.0275 | 0.0582 |  |  |
|  | Interaction | 0.0120 | 0.0625 | 0.0374 | 0.85 |
| *Combined*<br>*(no interaction)* | Intercept | 1.8069 | 0.0429 |  |  |
|  | Treatment_female | -0.0158 | 0.0299 | 0.288 | 0.59 |
|  | Origin_female | -0.0220 | 0.0506 | 0.187 | 0.67 |
| Log10 (latency) ~ treatment_male*origin_male + (1\|female_ID) + (1\|rival_pair) + (1\|site_male),<br>family = Gaussian, link = identity |  |  |  |  |  |
| *Combined*<br>*(interaction)* | Intercept | 1.7739 | 0.0487 |  |  |
|  | Treatment_male | 0.0226 | 0.0477 |  |  |
|  | Origin_male | -0.0235 | 0.0601 |  |  |
|  | Interaction | 0.0273 | 0.0615 | 0.196 | 0.66 |
| *Combined*<br>*(no interaction)* | Intercept | 1.7660 | 0.0453 |  |  |
|  | Treatment_male | 0.0389 | 0.0302 | 1.655 | 0.20 |
|  | Origin_male | -0.0102 | 0.0521 | 0.038 | 0.84 |
| cbind(choices deviating from equal choices, choices not deviating from equal choices) ~<br>treatment_female * origin_female + (1\|stimulus), family = binomial, link= logit |  |  |  |  |  |
| *Combined*<br>*(interaction)* | Intercept | 0.220 | 0.431 |  |  |
|  | Treatment_female | -0.666 | 0.562 |  |  |
|  | Origin_female | -1.166 | 0.486 |  |  |
|  | Interaction | 1.820 | 0.719 | 6.66 | **0.0099** |
| *Forest frogs* | Intercept | 0.220 | 0.431 |  |  |
|  | Treatment_female | -0.666 | 0.562 | 1.186 | 0.2358 |
| *Urban frogs* | Intercept | -0.946 | 0.363 |  |  |
|  | Treatment_female | 1.154 | 0.420 | -2.745 | **0.0061** |
| cbind(choices deviating from equal choices, choices not deviating from equal choices) ~<br>treatment_male * origin_male + (1\|rival_pair), family = binomial, link= logit |  |  |  |  |  |
| *Combined*<br>*(interaction)* | Intercept | -0.542 | 0.570 |  |  |
|  | Treatment_male | 1.028 | 0.474 |  |  |
|  | Origin_male | 0.415 | 0.756 |  |  |
|  | Interaction | -1.020 | 0.603 | 2.89 | **0.089** |
| *Combined*<br>*(no interaction)* | Intercept | -0.2311 | 0.5241 |  |  |
|  | Treatment_male | 0.4071 | 0.2899 | 1.972 | 0.16 |
|  | Origin_male | -0.0974 | 0.6733 | 0.021 | 0.88 |
| *Forest frogs* | Intercept | -0.542 | 0.570 |  |  |
|  | Treatment_male | 1.0276 | 0.474 | -2.166 | **0.0303** |
| *Urban frogs* | Intercept | -0.127 | 0.500 |  |  |
|  | Treatment_male | 0.0072 | 0.373 | -0.019 | 0.9845 |

**Figure S4.**
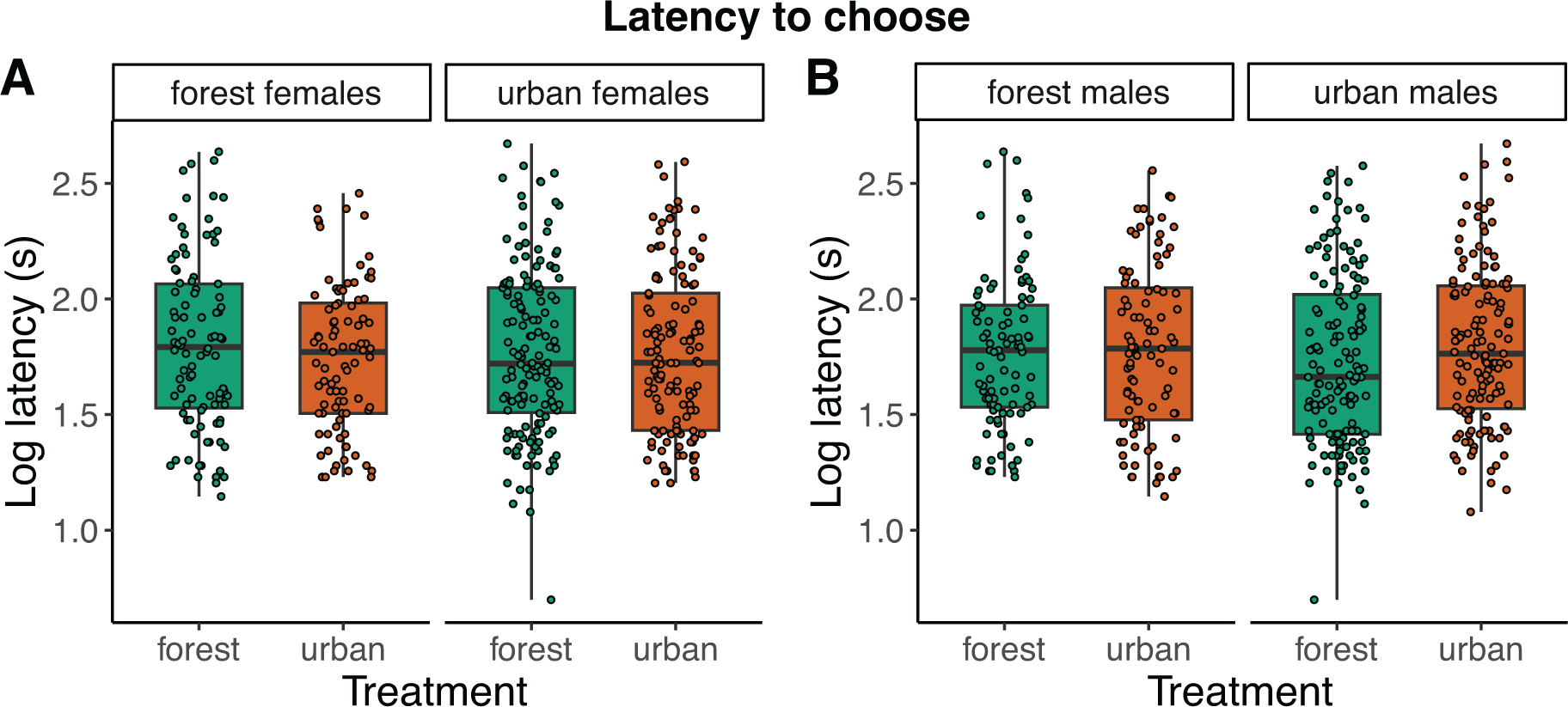
Latency to choose (log seconds) split in A) preference of urban and forest females choosing under urban or forest sensory conditions and split in B) preference for urban and forest rival pairs interacting under urban or forest sensory conditions. Graph depicts raw data. See main text for statistics.

**Table S7.** Preference strength (0-1) obtained in the phonotaxis experiments and 95% confidence interval (CI) based on 1000 simulations of random choices. Both unweighted and weighted (based on number of choices per stimulus or rival pairs) are reported. Data is split in female origin and female sensory treatment, or in male origin and male sensory treatment.

| Origin | Sensory treatment | Measured female preference (weighted) | Unweighted simulated preference strength (95% CI) | Weighted simulated preference strength (95% CI) |
| --- | --- | --- | --- | --- |
| **Female origin and sensory treatment** |  |  |  |  |
| *Forest females* | Forest | 0.61 (0.55) | 0.08; 0.48 | 0.08; 0.45 |
|  | Urban | 0.42 (0.42) | 0.10; 0.50 | 0.10; 0.48 |
| *Urban females* | Forest | 0.34 (0.32) | 0.11; 0.40 | 0.11; 0.39 |
|  | Urban | 0.55 (0.53) | 0.10; 0.42 | 0.11; 0.40 |
| **Male origin and sensory treatment** |  |  |  |  |
| *Forest males* | Forest | 0.35 (0.35) | 0.00; 0.38 | 0.00; 0.39 |
|  | Urban | 0.58 (0.57) | 0.00; 0.39 | 0.00; 0.38 |
| *Urban males* | Forest | 0.46 (0.46) | 0.03; 0.32 | 0.03; 0.32 |
|  | Urban | 0.47 (0.46) | 0.04; 0.33 | 0.04; 0.33 |

**Figure S5.**
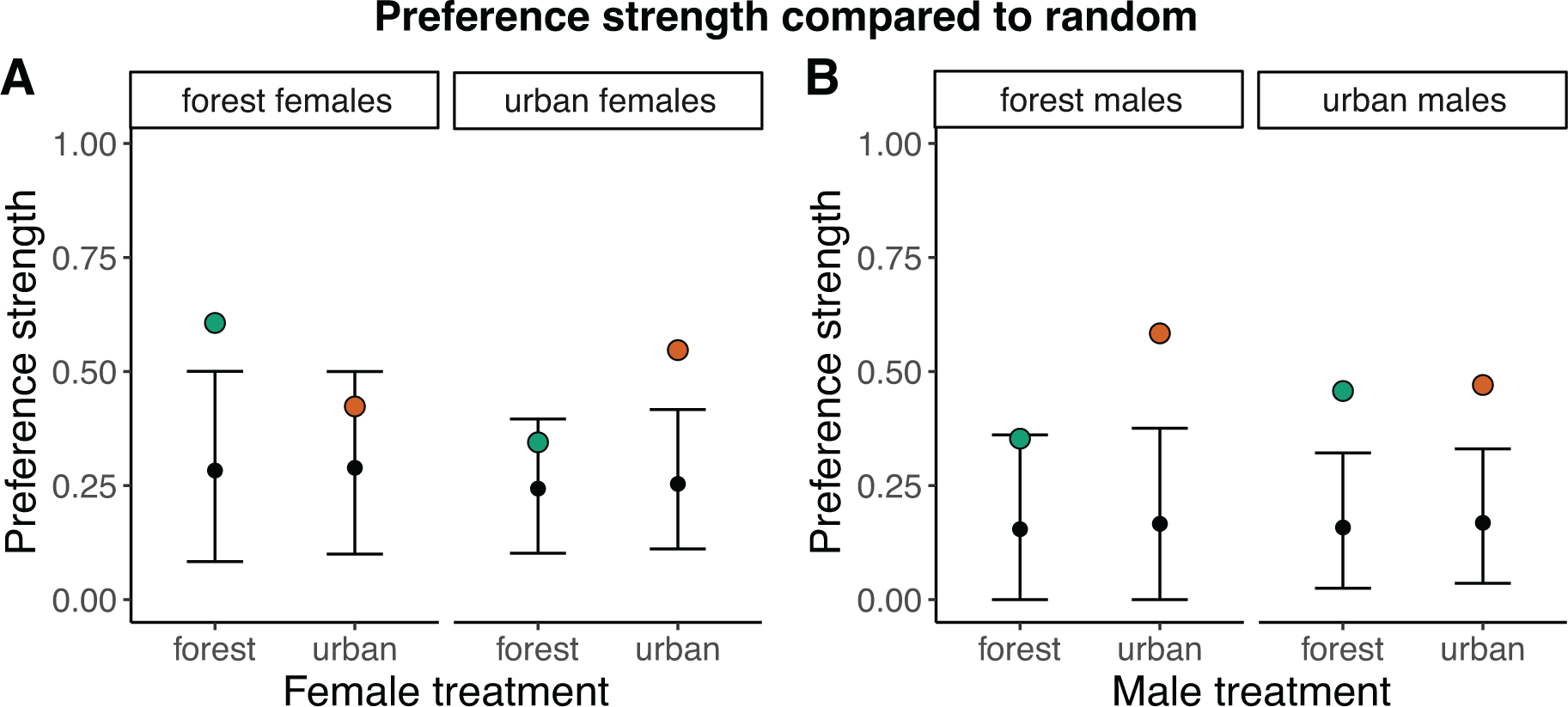
Preference strength (0-1) split in A) preference of urban and forest females choosing under urban or forest sensory conditions and split in B) preference for urban and forest rival pairs interacting under urban or forest sensory conditions. Open circles indicate preference strength obtained in the phonotaxis experiments and the error bars indicated 95% confidence intervals based on 1000 simulations. Mean preference strength was not weighted depending on the number of choices per stimulus or rival pair. See main text and Table X for statistics.

**Table S8.** Results on the association between preference strength and interaction level characteristics (proportional call rate difference, interaction length, proportional call complexity difference, overlap rate and proportional P2P whine amplitude difference). Table shows the result of weighted model averaging (delta < 4). See main text for details on statistics.

| Explanatory | Estimate | s.e. | z value | P |
| --- | --- | --- | --- | --- |
| cbind(choices deviating from equal choices, choices not deviating from equal choices) ~<br>prop_diff_call_rate + log10(interaction length) + prop_diff_complexity + overlap rate +<br>prop_diff_P2P, (1\|rival_pair), family = binomial, link= logit |  |  |  |  |
| Intercept | -0.16508011 | 0.2524651 |  |  |
| prop_diff_call_rate | 0.57309120 | 0.2201351 | 2.4803655 | **0.0131248** |
| log10(interaction length) | -0.51587032 | 0.2141835 | 2.2951737 | **0.0217232** |
| prop_diff_complexity | 0.23178010 | 0.1948456 | 1.1320449 | 0.2576156 |
| overlap rate | 0.14072933 | 0.2309802 | 0.5798130 | 0.5620407 |
| prop_diff_P2P | -0.06987971 | 0.2106951 | 0.3156274 | 0.7522853 |

## References

Akre KL, Farris HE, Lea AM, Page RA, Ryan MJ. 2011. Signal Perception in Frogs and Bats and the Evolution of Mating Signals. Science (80-). 333(6043):751–752. doi:10.1126/science.1205623. http://www.sciencemag.org/cgi/doi/10.1126/science.1205623.

Akre KL, Johnsen S. 2014. Psychophysics and the evolution of behavior. Trends Ecol Evol. 29(5):291–300. doi:10.1016/j.tree.2014.03.007.

Andersson M. 1994. Sexual Selection. Princeton: Princeton University press.

Arnott G, Elwood RW. 2008. Information gathering and decision making about resource value in animal contests. Anim Behav. 76(3):529–542. doi:10.1016/j.anbehav.2008.04.019.

Barber JR, Crooks KR, Fristrup KM. 2010. The costs of chronic noise exposure for terrestrial organisms. Trends Ecol Evol. 25(3):180–189. doi:10.1016/j.tree.2009.08.002.

Bartoń K. 2023. MuMIn: multi-model inference. https://cran.r-project.org/package=MuMIn.

Bates D, Mächler M, Bolker B, Walker S. 2015. Fitting Linear Mixed-Effects Models Using lme4. J Stat Softw. 67(1). doi:10.18637/jss.v067.i01. http://www.jstatsoft.org/v67/i01/.

Bateson M, Healy SD. 2005. Comparative evaluation and its implications for mate choice. Trends Ecol Evol. 20(12):659–664. doi:10.1016/j.tree.2005.08.013.

Bernal XE, Akre KL, Baugh AT, Rand AS, Ryan MJ. 2009. Female and male behavioral response to advertisement calls of graded complexity in túngara frogs, Physalaemus pustulosus. Behav Ecol Sociobiol. 63(9):1269–1279. doi:10.1007/s00265-009-0795-5.

Bernal XE, Page RA, Rand AS, Ryan MJ. 2007. Cues for eavesdroppers: Do frog calls indicate prey density and quality? Am Nat. 169(3):409–415. doi:10.1086/510729.

Bernal XE, Rand a. S, Ryan MJ. 2006. Acoustic preferences and localization performance of blood-sucking flies (Corethrella Coquillett) to túngara frog calls. Behav Ecol. 17(5):709–715. doi:10.1093/beheco/arl003. http://academic.oup.com/beheco/article/17/5/709/206929/Acoustic-preferences-and-localization-performance.

Bird S, Parker J. 2014. Low levels of light pollution may block the ability of male glow-worms (Lampyris noctiluca L.) to locate females. J Insect Conserv. 18(4):737–743. doi:10.1007/s10841-014-9664−2.

Bonachea LA, Ryan MJ. 2011. Simulated Predation Risk Influences Female Choice in Túngara Frogs, Physalaemus pustulosus. Ethology. 117(5):400–407. doi:10.1111/j.1439-0310.2011.01889.x.

Brumm H, Slabbekoorn H. 2005. Acoustic Communication in Noise. Adv Study Behav. 35(05):151–209. doi:10.1016/S0065-3454(05)35004−2.

Callander S, Hayes CL, Jennions MD, Backwell PRY. 2013. Experimental evidence that immediate neighbors affect male attractiveness. Behav Ecol. 24(3):730–733. doi:10.1093/beheco/ars208.

Calsbeek R, Zamora-Camacho FJ, Symes LB. 2022. Individual contributions to group chorus dynamics influence access to mating opportunities in wood frogs. Ecol Lett. 25(6):1401–1409. doi:10.1111/ele.14002.

Candolin U, Wong BBM. 2019. Mate choice in a polluted world: Consequences for individuals, populations and communities. Philos Trans R Soc B Biol Sci. 374(1781). doi:10.1098/rstb.2018.0055.

Coss DA, Hunter KL, Taylor RC. 2020. Silence is sexy: soundscape complexity alters mate choice in túngara frogs. Behav Ecol.:1–11. doi:10.1093/beheco/araa091.

Costello RA, Symes LB. 2014. Effects of anthropogenic noise on male signalling behaviour and female phonotaxis in Oecanthus tree crickets. Anim Behav. 95:15–22. doi:10.1016/j.anbehav.2014.05.009. 10.1016/j.anbehav.2014.05.009.

Cronin AD, Smit JAH, Halfwerk W. 2022. Anthropogenic noise and light alter temporal but not spatial breeding behavior in a wild frog. Behav Ecol. 33(6):1115–1122. doi:10.1093/beheco/arac077.

Cronin AD, Smit JAH, Muñoz MI, Poirier A, Moran PA, Jerem P, Halfwerk W. 2022. A comprehensive overview of the effects of urbanisation on sexual selection and sexual traits. Biol Rev. 97(4):1325–1345. doi:10.1111/brv.12845. https://onlinelibrary.wiley.com/doi/10.1111/brv.12845.

Cynx J, Lewis R, Tavel B, Tse H. 1998. Amplitude regulation of vocalizations in noise by a songbird, Taeniopygia guttata. Anim Behav. 56(1):107–113. doi:10.1006/anbe.1998.0746.

Darwin CR. 1871. The Descent of Man, and Selection in Relation to Sex. London, UK: John Murray.

Dominoni DM, Halfwerk W, Baird E, Buxton RT, Fernández-Juricic E, Fristrup KM, McKenna MF, Mennitt DJ, Perkin EK, Seymoure BM, et al. 2020. Why conservation biology can benefit from sensory ecology. Nat Ecol Evol. 4(4):502–511. doi:10.1038/s41559-020-1135-4. http://www.nature.com/articles/s41559-020-1135-4.

Francis CD, Ortega CP, Cruz A. 2009. Noise Pollution Changes Avian Communities and Species Interactions. Curr Biol. 19(16):1415–1419. doi:10.1016/j.cub.2009.06.052. 10.1016/j.cub.2009.06.052.

van Geffen KG, van Eck E, de Boer RA, van Grunsven RHA, Salis L, Berendse F, Veenendaal EM. 2015. Artificial light at night inhibits mating in a Geometrid moth. Stewart A, Sait S, editors. Insect Conserv Divers. 8(3):282–287. doi:10.1111/icad.12116. https://onlinelibrary.wiley.com/doi/10.1111/icad.12116.

van Geffen KG, Groot AT, van Grunsven RHA, Donners M, Berendse F, Veenendaal EM. 2015. Artificial night lighting disrupts sex pheromone in a noctuid moth. Ecol Entomol. 40(4):401–408. doi:10.1111/een.12202. https://onlinelibrary.wiley.com/doi/10.1111/een.12202.

Gerhardt HC, Huber F. 2002. Acoustic communication in insects and anurans: Common problems and diverse solutions. Chicago: University of Chicago Press.

Gerhardt HC, Roberts JD, Bee MA, Schwartz JJ. 2000. Call matching in the quacking frog (Crinia georgiana). Behav Ecol Sociobiol. 48(3):243–251. doi:10.1007/s002650000226. http://link.springer.com/10.1007/s002650000226.

Goutte S, Kime NM, Argo IV TF, Ryan MJ. 2010. Calling strategies of male túngara frogs in response to dynamic playback. Behaviour. 147(1):65–83. doi:10.1163/000579509X12483520922205. https://brill.com/view/journals/beh/147/1/article-p65_5.xml.

Green AJ. 1990. Determinants of chorus participation and the effects of size, weight and competition on advertisement calling in the tungara frog, Physalaemus pustulosus (Leptodactylidae). Anim Behav. 39(4):620– 638. doi:10.1016/S0003-3472(05)80373−2.

Greenfield MD. 2005. Mechanisms and Evolution of Communal Sexual Displays in Arthropods and Anurans. In: Advances in the Study of Behavior. Vol. 35. p. 1–62. http://linkinghub.elsevier.com/retrieve/pii/S0065345405350017.

Greenfield MD. 2015. Signal interactions and interference in insect choruses: singing and listening in the social environment. J Comp Physiol A Neuroethol Sensory, Neural, Behav Physiol. 201(1):143–154. doi:10.1007/s00359-014-0938-7.

Greenfield MD, Rand AS. 2000. Frogs Have Rules: Selective Attention Algorithms Regulate Chorusing in Physalaemus pustulosus (Leptodactylidae). Ethology. 106(4):331–347. doi:10.1046/j.1439-0310.2000.00525.x. http://doi.wiley.com/10.1046/j.1439-0310.2000.00525.x.

Halfwerk W, Blaas M, Kramer L, Hijner N, Trillo PA, Bernal XE, Page RA, Goutte S, Ryan MJ, Ellers J. 2019. Adaptive changes in sexual signalling in response to urbanization. Nat Ecol Evol. 3(3):374–380. doi:10.1038/s41559-018-0751-8. 10.1038/s41559-018-0751-8.

Halfwerk W, Bot S, Buikx J, van der Velde M, Komdeur J, ten Cate C, Slabbekoorn H. 2011. Low-frequency songs lose their potency in noisy urban conditions. Proc Natl Acad Sci. 108(35):14549–14554. doi:10.1073/pnas.1109091108. http://www.pnas.org/cgi/doi/10.1073/pnas.1109091108.

Halfwerk W, Lea a. M, Guerra M a., Page R a., Ryan MJ. 2016. Vocal responses to noise reveal the presence of the Lombard effect in a frog. Behav Ecol. 27(2):669–676. doi:10.1093/beheco/arv204.

Halfwerk W, Slabbekoorn H. 2015. Pollution going multimodal: the complex impact of the human-altered sensory environment on animal perception and performance. Biol Lett. 11(4):20141051. doi:10.1098/rsbl.2014.1051. http://rsbl.royalsocietypublishing.org/content/11/4/20141051.article-info.

Harding HR, Gordon TAC, Eastcott E, Simpson SD, Radford AN. 2019. Causes and consequences of intraspecific variation in animal responses to anthropogenic noise. Behav Ecol. 30(6):1501–1511. doi:10.1093/beheco/arz114.

Hartig F. 2022. DHARMa: Residual Diagnostics for Hierarchical (Multi-Level / Mixed) Regression Models. https://cran.r-project.org/package=DHARMa.

Huet des Aunay G, Slabbekoorn H, Nagle L, Passas F, Nicolas P, Draganoiu TI. 2014. Urban noise undermines female sexual preferences for low-frequency songs in domestic canaries. Anim Behav. 87(C):67–75. doi:10.1016/j.anbehav.2013.10.010. 10.1016/j.anbehav.2013.10.010.

Iglesias-Carrasco M, Head ML, Jennions MD, Cabido C. 2017. Secondary compounds from exotic tree plantations change female mating preferences in the palmate newt (Lissotriton helveticus). J Evol Biol. 30(10):1788–1795. doi:10.1111/jeb.13091.

Kempenaers B, Borgström P, Loës P, Schlicht E, Valcu M. 2010. Artificial night lighting affects dawn song, extra-pair siring success, and lay date in songbirds. Curr Biol. 20(19):1735–1739. doi:10.1016/j.cub.2010.08.028.

Kunc HP, Morrison K, Schmidt R. 2022. A meta-analysis on the evolution of the Lombard effect reveals that amplitude adjustments are a widespread vertebrate mechanism. Proc Natl Acad Sci U S A. 119(30):1–7. doi:10.1073/pnas.2117809119.

Kuriwada T. 2023. Differences in male calling song and female mate location behaviour between urban and rural crickets. Biol J Linn Soc. 139(3):275–285. doi:10.1093/biolinnean/blad027. https://academic.oup.com/biolinnean/article/139/3/275/7192923.

Kurvers RHJM, Hölker F. 2015. Bright nights and social interactions: A neglected issue. Behav Ecol. 26(2):334–339. doi:10.1093/beheco/aru223.

Kyba CCM, Kuester T, De Miguel AS, Baugh K, Jechow A, Hölker F, Bennie J, Elvidge CD, Gaston KJ, Guanter L. 2017. Artificially lit surface of Earth at night increasing in radiance and extent. Sci Adv. 3(11):1–9. doi:10.1126/sciadv.1701528.

Lampe U, Schmoll T, Franzke A, Reinhold K. 2012. Staying tuned: Grasshoppers from noisy roadside habitats produce courtship signals with elevated frequency components. Funct Ecol. 26(6):1348–1354. doi:10.1111/1365−2435.12000.

Larter LC, Bernal XE, Page RA, Ryan MJ. 2022. Local competitive environment and male condition influence within-bout calling patterns in túngara frogs. Bioacoustics. 32(2):121–142. doi:10.1080/09524622.2022.2070544. 10.1080/09524622.2022.2070544.

Larter LC, Ryan MJ. 2024. Female Preferences for More Elaborate Signals Are an Emergent Outcome of Male Chorusing Interactions in Túngara Frogs. Am Nat. 203(1). doi:10.1086/727469. https://www.journals.uchicago.edu/doi/10.1086/727469.

LaZerte SE, Slabbekoorn H, Otter KA. 2017. Territorial black-capped chickadee males respond faster to high-than to low-frequency songs in experimentally elevated noise conditions. PeerJ. 2017(4):1–19. doi:10.7717/peerj.3257.

Lea AM, Ryan MJ. 2015. Irrationality in mate choice revealed by tungara frogs. Science (80-). 349(6251):964– 966. doi:10.1126/science.aab2012. http://www.sciencemag.org/content/349/6251/964.abstract%5Cn http://www.sciencemag.org/cgi/doi/10.1126/science.aab2012.

Lenth R V. 2023. emmeans: Estimated Marginal Means, aka Least-Squares Means. https://cran.r-project.org/package=emmeans.

Lowry H, Lill A, Wong BBM. 2013. Behavioural responses of wildlife to urban environments. Biol Rev. 88(3):537–549. doi:10.1111/brv.12012.

Luther D, Magnotti J. 2014. Can animals detect differences in vocalizations adjusted for anthropogenic noise? Anim Behav. 92:111–116. doi:10.1016/j.anbehav.2014.03.033. 10.1016/j.anbehav.2014.03.033.

McGregor PK, Peake TM. 2000. Communication networks: social environments for receiving and signalling behaviour. Acta Ethol. 2(2):71–81. doi:10.1007/s102110000015. http://academic.oup.com/hwj/article/17/1/3/627575.

McMahon TA, Rohr JR, Bernal XE. 2017. Light and noise pollution interact to disrupt interspecific interactions. Ecology. 98(5):1290–1299. doi:10.1002/ecy.1770. https://onlinelibrary.wiley.com/doi/10.1002/ecy.1770.

Mennill DJ, Ratcliffe LM, Boag PT. 2002. Female eavesdropping on male song contests in songbirds. Science (80-). 296(5569):873. doi:10.1126/science.296.5569.873.

Møller AP. 2010. Interspecific variation in fear responses predicts urbanization in birds. Behav Ecol. 21(2):365–371. doi:10.1093/beheco/arp199.

Naguib M. 2013. Living in a noisy world: Indirect effects of noise on animal communication. Behaviour. 150(9–10):1069–1084. doi:10.1163/1568539X-00003058.

Neelon DP, Höbel G. 2019. Staying ahead of the game—plasticity in chorusing behavior allows males to remain attractive in different social environments. Behav Ecol Sociobiol. 73(9). doi:10.1007/s00265-019−2737-1.

Otter KA, Ratcliffe L, Njegovan M, Fotheringham J. 2002. Importance of frequency and temporal song matching in black-capped chickadees: Evidence from interactive playback. Ethology. 108(2):181–191. doi:10.1046/j.1439-0310.2002.00764.x.

Rand AS, Bridarolli ME, Dries L, Ryan MJ. 1997. Light Levels Influence Female Choice in Túngara Frogs: Predation Risk Assessment? Copeia. 1997(2):447. doi:10.2307/1447770. https://www.jstor.org/stable/1447770?origin=crossref.

Rand AS, Ryan MJ. 1981. The adaptive significance of a complex vocal repertoire in a Neotropical frog. Z Tierpsychol. 57(3–4):209–214. doi:10.1111/j.1439-0310.1981.tb01923.x.

Ryan MJ. 1980. Female mate choice in a Neotropical frog. Science (80-). 209(4455):523–525. doi:10.1126/science.209.4455.523. http://www.ncbi.nlm.nih.gov/pubmed/17831371%5Cn http://www.sbs.utexas.edu/ryan/Publications/1980-82/1980Science209-523.pdf.

Ryan MJ. 1985. The túngara frog. Chicago: The University of Chicago Press.

Ryan MJ, Tuttle MD, Rand AS. 1982. Bat predation and sexual advertisement in a Neotropical anuran. Am Nat. 199(1):136–139.

Slabbekoorn H, Peet M. 2003. Birds sing at a higher pitch in urban noise. Nature. 424(6946):267–267. doi:10.1038/424267a. http://www.nature.com/articles/424267a.

Smit JAH, Cronin AD, van der Wiel I, Oteman B, Ellers J, Halfwerk W. 2022. Interactive and independent effects of light and noise pollution on sexual signaling in frogs. Front Ecol Evol. 10(August):1–12. doi:10.3389/fevo.2022.934661. https://www.frontiersin.org/articles/10.3389/fevo.2022.934661/full.

Smit JAH, Vooijs R, Lindenburg P, Baugh AT, Halfwerk W. 2024. Noise and light pollution elicit endocrine responses in urban but not forest frogs. Horm Behav. 157(November 2023):105453. doi:10.1016/j.yhbeh.2023.105453. 10.1016/j.yhbeh.2023.105453.

Sol D, Lapiedra O, González-Lagos C. 2013. Behavioural adjustments for a life in the city. Anim Behav. 85(5):1101–1112. doi:10.1016/j.anbehav.2013.01.023. 10.1016/j.anbehav.2013.01.023.

Sueur J, Aubin T, Simonis C. 2008. seewave: a free modular tool for sound analysis and synthesis. Bioacoustics, 18: 213–226. Bioacoustics. 18:213–226.

Tárano Z. 2015. Choosing a Mate in a Cocktail Party-Like Situation: The Effect of Call Complexity and Call Timing between Two Rival Males on Female Mating Preferences in the Túngara Frog Physalaemus pustulosus. Ethology. 121(8):749–759. doi:10.1111/eth.12387.

Taylor RC, Wilhite KO, Ludovici RJ, Mitchell KM, Halfwerk W, Page RA, Ryan MJ, Hunter KL. 2021. Complex sensory environments alter mate choice outcomes. J Exp Biol. 224(1). doi:10.1242/jeb.233288.

Team R Development Core. 2022. A Language and Environment for Statistical Computing. R Found Stat Comput. 2:https://www.R-project.org. http://www.r-project.org.

Tuttle MD, Taft LK, Ryan MJ. 1982. Evasive behaviour of a frog in response to bat predation. Anim Behav. 30(2):393–397. doi:10.1016/S0003-3472(82)80050-X. https://linkinghub.elsevier.com/retrieve/pii/S000334728280050X.

Uchida K, Suzuki KK, Shimamoto T, Yanagawa H, Koizumi I. 2019. Decreased vigilance or habituation to humans? Mechanisms on increased boldness in urban animals. Behav Ecol.:1–8. doi:10.1093/beheco/arz117.

Weber EH. 1948. The sense of touch and common feeling, 1846. In: Readings in the history of psychology. East Norwalk: Appleton-Century-Crofts. p. 194–196. http://content.apa.org/books/11304-023.

Wickham H. 2016. ggplot2: Elegant graphics for data analysis. Springer-Verlag New York. https://ggplot2.tidyverse.org.

Wiley RH, Richards DG. 1978. Physical constraints on acoustic communication in the atmosphere: Implications for the evolution of animal vocalizations. Behav Ecol Sociobiol. 3(1):69–94. doi:10.1007/BF00300047. http://link.springer.com/10.1007/BF00300047.

Zuur AF, Ieno EN, Elphick CS. 2010. A protocol for data exploration to avoid common statistical problems. Methods Ecol Evol. 1(1):3–14. doi:10.1111/j.2041−210x.2009.00001.x.

